# Single-cell atlas of developing mouse palates reveals cellular and molecular transitions in periderm cell fate

**DOI:** 10.1101/2024.10.28.620753

**Authors:** Wenbin Huang, Zhenwei Qian, Jieni Zhang, Yi Ding, Bin Wang, Jiuxiang Lin, Xiannian Zhang, Huaxiang Zhao, Feng Chen

**Affiliations:** Department of Orthodontics, Peking University School and Hospital of Stomatology, Beijing 100081, China; Key Laboratory of Shaanxi Province for Craniofacial Precision Medicine Research, College of Stomatology, Xi’an Jiaotong University, Xi’an 710004, China; Guangdong Provincial High-level Clinical Key Specialty, Guangdong Province Engineering Research Center of Oral Disease Diagnosis and Treatment, Department of Orthodontics, Stomatological Center, Peking University Shenzhen Hospital, Shenzhen Peking University-The Hong Kong University of Science and Technology Medical Center, Shenzhen 518036, China; Department of Materials Science and Engineering, Southern University of Science and Technology, Shenzhen 518055, China; Peking University 302 Clinical Medical School, Beijing 100039, China; Department of Neurobiology, School of Basic Medical Sciences, Beijing Key Laboratory of Neural Regeneration and Repair, Capital Medical University, Beijing 100069, China; National Center of Stomatology, National Clinical Research Center for Oral Diseases, National Engineering Laboratory for Digital and Material Technology of Stomatology, Beijing Key Laboratory for Digital Stomatology, Research Center of Engineering and Technology for Computerized Dentistry Ministry of Health, NMPA Key Laboratory for Dental Materials, Beijing 100081, China; Department of Physiology and Pathophysiology, School of Basic Medical Sciences, Xi’an Jiaotong University, Xi’an 710049, China; Department of Special Clinic, College of Stomatology, Xi’an Jiaotong University, Xi’an 710004, China; Department of Orthodontics, College of Stomatology, Xi’an Jiaotong University, Xi’an 710004, China; Central laboratory, Peking University School and Hospital of Stomatology, Beijing 100081, China

**Keywords:** Cleft palate, Single-cell sequencing, Palatogenesis, Periderm, Cell fate

## Abstract

Cleft palate is one of the most common congenital craniofacial disorders that affects children’s appearance and oral functions. Investigating the transcriptomics during palatogenesis is crucial for comprehending the etiology of this disorder and facilitating prenatal molecular diagnosis. However, there is limited knowledge about the single-cell differentiation dynamics during mid- and late-palatogenesis, specifically regarding the subpopulations and developmental trajectories of periderm, a rare but critical cell population. Here we explore the single-cell landscape of mouse developing palates from E10.5 to E16.5. We systematically depict the single-cell transcriptomics of mesenchymal and epithelial cells during palatogenesis, including subpopulations and differentiation dynamics. Additionally, we identify four subclusters of palatal periderm and construct two distinct trajectories of cell fates for periderm cells. Our findings reveal that *Claudins* and *Arhgap29* play a role in the non-stick function of the periderm before the palatal shelves contact, and *Pitx2* mediates the adhesion of periderm during the contact of opposing palatal shelves. Furthermore, we demonstrate that epithelial-mesenchymal transition (EMT), apoptosis, and migration collectively contribute to the degeneration of periderm cells in the medial epithelial seam. Taken together, our study suggests a novel model of periderm development during palatogenesis and delineates the cellular and molecular transitions in periderm cell determination.

## Introduction

Orofacial clefts (OFCs) are the most prevalent congenital craniofacial disorders, affecting approximately 1/700 and 1/500 of live births worldwide and in China, respectively [1, 2]. These malformations affect not only the appearance but also the oral functions and mental health of children [3]. Cleft palates are one of the three types of OFCs, which can occur alone or in combination with cleft lips. Additionally, due to the alveolar bone obstructing the imaging, prenatal ultrasound may have difficulty in detecting cleft palates [4].

Palatogenesis is a conserved process in both humans and mice, and its disruption by genetic or environmental factors results in cleft palates [5]. In mice, palatogenesis begins at embryonic day (E) 10.5 and ends at around E16.0. The primary palate initiates at E10.5, followed by the paired secondary palatal shelves originating from the maxillary processes at E11.5. The palatal shelves start to grow vertically by E13.5 and as the tongue depresses at around E14.0, they quickly elevate and grow horizontally. Subsequently, the shelves contact and fuse, forming the intact roof of the oral cavity by approximately E16.0 [5, 6].

Apart from the various other cell types such as epithelial, mesenchymal, neuronal, and endothelial cells, periderm cells, a continuous single-cell layer of flattened cells, also play a critical role in palatogenesis [7, 8]. The oral periderm initially appears over the developing facial prominence and then covers the developing palatal shelves. Before the adhesion of the horizontally developing palatal shelves, periderm cells serve as a protective “non-stick” coat, preventing pathological adhesion between the palatal shelves and tongue, maxillary, or mandibular tissue [7–9]. However, once the opposing palatal shelves come into contact with each other, periderm might mediate the subsequent adhesion of palatal shelves [10, 11]. During the final phase of palatogenesis, the periderm in the medial edge epithelium might degenerate or migrate out of the region of fusion, ensuring the successful fusion of the palatal shelves [10]. The shifting functions of periderm, transitioning from preventing pathological adhesion to becoming a prerequisite for the fusion of palatal shelves, suggest the existence of different subtypes of periderm cells in palatogenesis [7, 8]. However, little is known about the developmental trajectories of these different subtypes of periderm cells and the key molecules that drive their development.

Recent advancements in high-resolution single-cell RNA sequencing (scRNA-seq) technology have provided a more detailed understanding of differentiation trajectories and cell fate determinations during embryogenesis [12, 13]. Although a previous study using scRNA-seq has explored the cellular and molecular transitions in the fusion region of the upper lip and primary palate during the early phase of palatogenesis in mice (E11.5) [14], our understanding of single-cell differentiation dynamics of the secondary palate during mid- and late-stages of palatogenesis (E13.5-E16.0) remains limited. In this study, we utilized scRNA-seq to probe into the single-cell landscape of mouse developmental palates from E10.5 to E16.5. Our results provide a more comprehensive view of the molecular events underpinning mouse palatogenesis. Specifically, we identified two developmental trajectories of periderm and four periderm subpopulations covering palatal shelves. We also uncovered key molecules that determine periderm cell fates and subsequently validated these findings.

## Results

### Single-cell RNA sequencing identifies 29 cell clusters during palatogenesis in mice

To provide a thorough view of mouse palatal development across all stages, we conducted scRNA-seq using the 10x Chromium platform on palatal shelves collected at four critical time points during mouse palatogenesis [6]: initial stage (E10.5), vertical growth stage (E13.5), adherence and fusion initiation stage (E15.0) and completion stage (E16.5). Overall, we obtained an average of 3,359 genes and 14,633 counts from 41,419 cells (12,368 in E10.5, 8,950 in E13.5, 9,173 in E15.0, and 10,928 in E16.5) with high sequencing saturation and a high ratio of reads mapping to the genome (**Figure 1**A, Figure S1, Figure S2).

**Figure 1.**
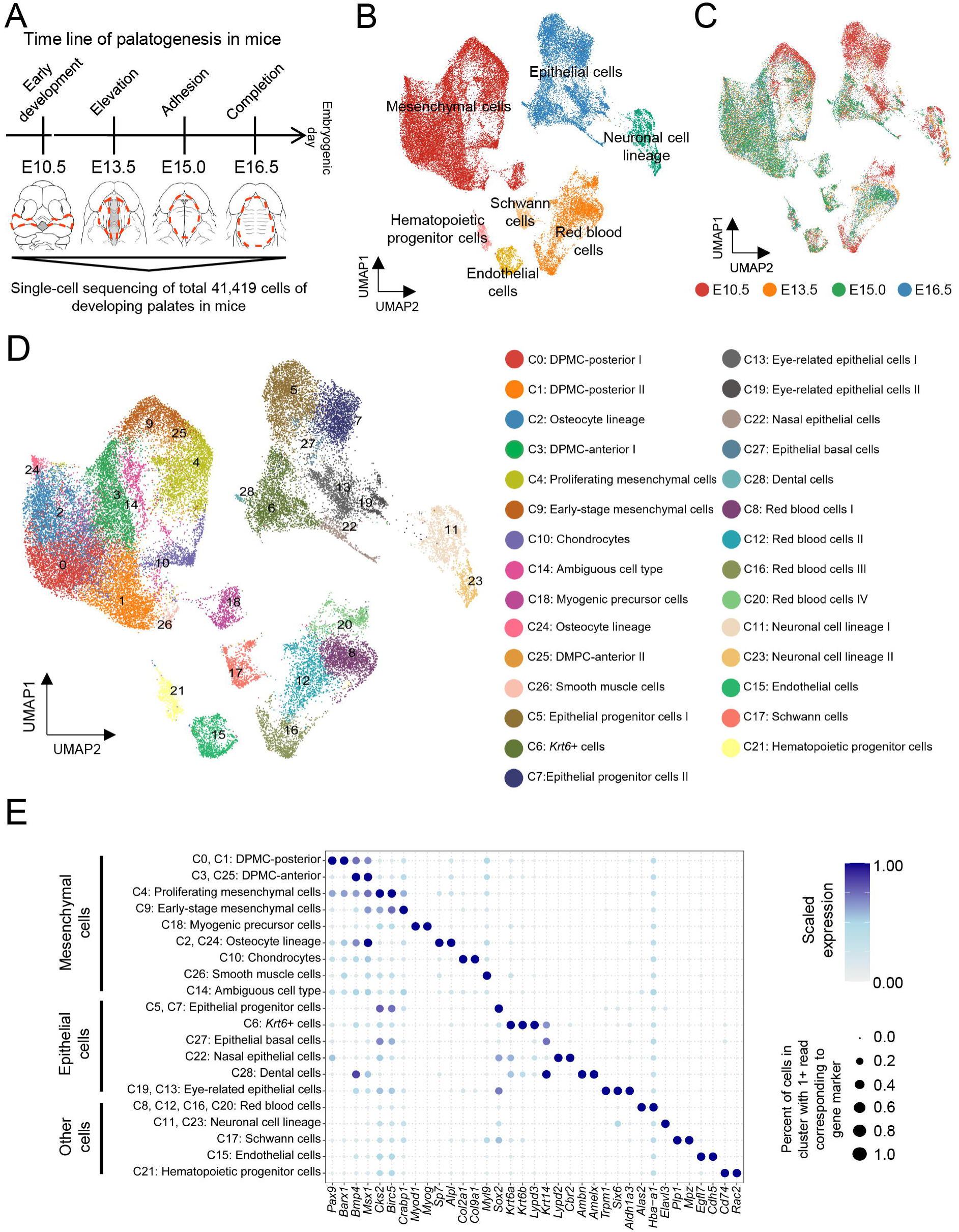
Identification of 29 cell clusters and 7 major cell types through scRNA-seq during mouse palatogenesis. **(A)** Schematic diagram illustrating the tissue samples sequenced at four developmental stages (E10.5 to E16.5) during mouse palatogenesis. The dashed circles indicate the regions of isolated tissue. For more sequencing details, refer to Figure S1. **(B-D)** UMAP plot of 41,419 cells visualizes general structure colored by **(B)** 7 major cell types, **(C)** 4 time points, and **(D)** 29 cell clusters (C0-C28). **(E)** Dot plot showing the relative expression of selected marker genes per cell cluster. The dot size represents the percentage of cells within a cell cluster where the relevant markers were detected, and the color indicates the average expression level. ScRNA-seq, single-cell RNA sequencing.

With dimensionality reduction, 29 distinct cell clusters (C0-C28), which could be grouped into 7 major cell types, were visualized in UMAP analysis. Based on the expression of *Pax9*, *Col3a1*, *Lypd3*, *Trp63*, *Alas2*, *Elav3*, *Plp1*, *Egfl7*, and *Rac2*, we found that 12 clusters exhibited a mesenchymal profile (C0-C4, C9, C10, C14, C18, C24-C26), 8 clusters exhibited an epithelial profile (C5-C7, C13, C19, C22, C27, C28), 4 clusters had a profile of red blood cells (C8, C12, C16, C20), 2 clusters exhibited a neuronal profile (C11, C23), while C17 showed a profile of Schwann cells, C15 exhibited an endothelial profile, and C21 displayed a profile of hematopoietic progenitor cells (Figure 1B-E, Figure S3, Figure S4; Table S1). To validate the existence of mesenchymal and epithelial populations, we also performed immunofluorescence (IF) assays using marker genes on coronal sections of mouse heads at E10.5, E13.5, E15.0, and E16.5 (COL3A1 for the mesenchymal population [15] and TRP63 for the epithelial population [16]), which the results were consistent with our scRNA-seq data (Figure S5A and B). The emergence of these cell clusters was associated with the progression along the sample time points. At the onset of palatogenesis starting from E10.5, the cell clusters gradually diversified (Figure S6A), consistent with previous studies [14]. Additionally, the analysis of sample distribution across different time points within each cluster revealed that cells belonging to C5, C7, C20, C13, and C9 exhibited a relatively early developmental phenotype, whereas cells in C26 and C28 showed a more mature phenotype (Figure S6B).

### Subpopulations and dynamic differentiation of mesenchymal cells

During mouse palatogenesis, mesenchymal and epithelial cells were the most dominant cell types, apart from the cells that form blood vessels and nerves, which exhibited a more complex architecture (Figure 1, Figure S3, Figure S4). Among the 12 clusters of mesenchymal cells, C2 and C24 exhibited characteristics of osteocytes with high and specific expression of *Sp7* and *Alpl* [17, 18]. Additionally, C10 displayed characteristics of chondrocytes with specific expression of *Col2a1* and *Col9a1* [19, 20]. C18 and C26 expressed markers associated with muscle formation, with C18 expressing muscle progenitor markers (*Myod1* and *Myog*) [21] and C26 showing a more differentiated state and identified as smooth muscle cells [22]. Based on the longitudinal gene expression pattern in the mouse developing palate [6, 23], we identified C3 and C25 as the developing palatal mesenchymal cells (DPMCs) in the anterior region, and C0 and C1 as the DPMCs in the posterior region. Moreover, C4 expressed critical genes for early proliferation [24, 25], suggesting that these cells represented the proliferating mesenchymal cells, and C9 expressed high levels of the pluripotent neural crest cell marker *Crabp1* [26], annotating them as early-stage mesenchymal cells. However, C14, located in the most central position of the UMAP plot of mesenchymal cells, did not significantly express any anatomy-/lineage-specific genes in heatmap analysis and therefore was ambiguous in nature (Figure 1D and E, Figure S3, Figure S4; Table S1).

To better understand the developmental progressions that occur in mesenchyme during palatogenesis in mice, we conducted pseudotime analysis on these 12 mesenchymal cell clusters (**Figure 2**A). Based on data gleaned from samples taken at each time point, we identified C4 and C9 as the initiating cell populations that drove mesenchymal developmental dynamics and gave rise to two distinct lineages (Figure 2A-C, Figure S7). In addition, we identified 473 highly dispersed differentially expressed (DE) genes that could be classified into 5 distinct stages across pseudotime (Stage 1 to 5 in Figure 2D). The earliest stage was characterized by the enrichment of genes involved in cell proliferation preparation, including chromosome segregation and nuclear division. As pseudotime progressed, the mesenchyme started to proliferate and differentiate into neural crest cells and smooth muscle tissues. During Stage 3, a group of genes, including *Dlx5* and *Tagln2*, emerged, indicating that the muscle tissue had started to contract. In the last two stages, we observed gene sets involved in cartilage development, bone morphogenesis and extracellular matrix organization, which suggested that these mesenchymal populations had switched to mineralization and were forming a mineralized matrix (Figure 2D; Table S2).

**Figure 2.**
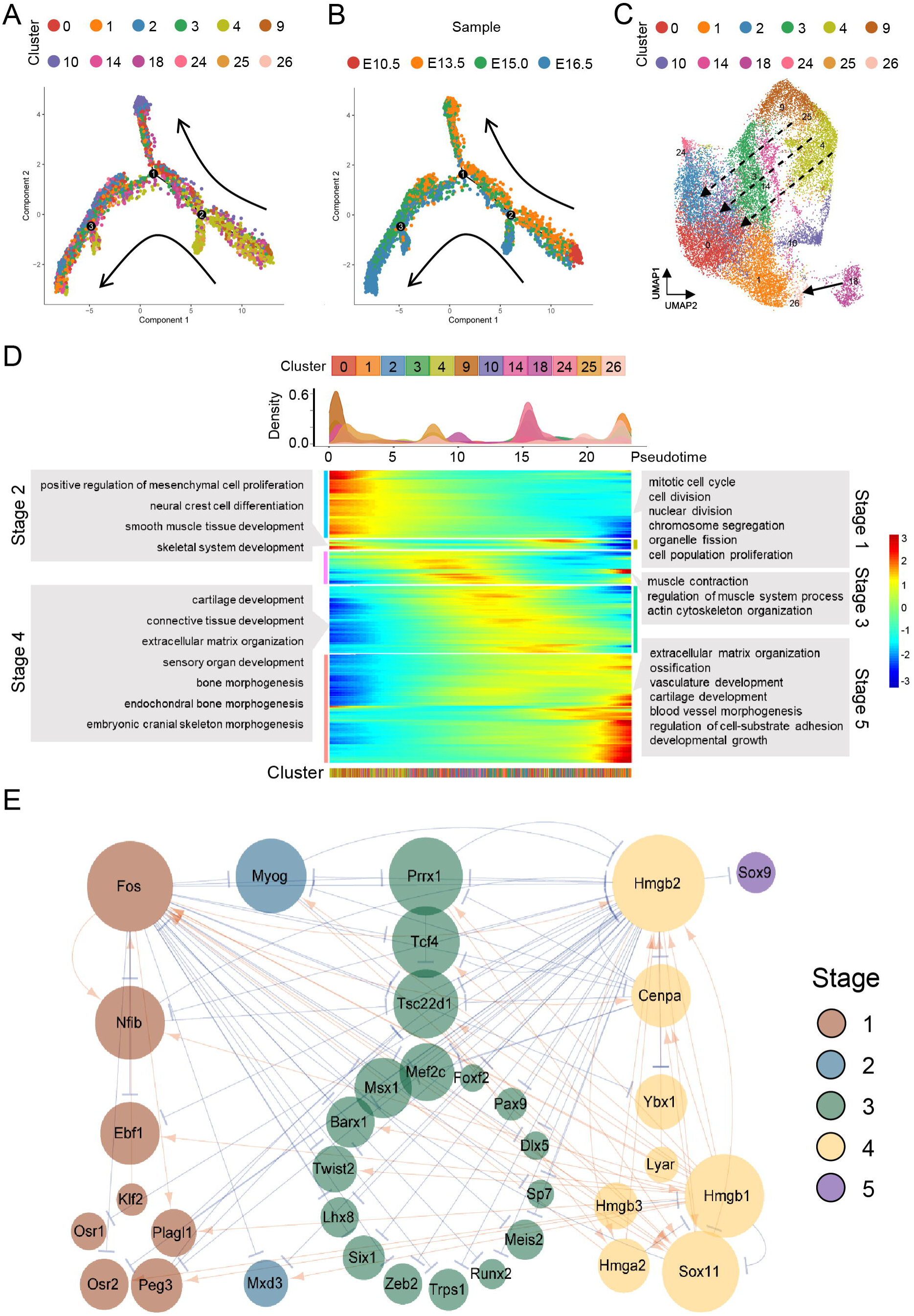
Characterization of dynamic differentiation of mesenchymal cells during palatogenesis in mice. **(A and B)** Pseudotime reconstruction of mesenchymal development reveals a bifurcation ordering of cells. The cells are colored by **(A)** clusters and **(B)** time points. Arrows indicate the major directions of developmental trajectories. **(C)** UMAP visualization of mesenchymal trajectories colored by cell clusters. Dotted arrows indicate the main developmental direction, while the arrow represents the trajectory from myogenic precursor cells to smooth muscle cells. **(D)** Expression heatmap of 473 dynamically expressed genes ordered across pseudotime, illustrating mesenchymal differentiation occurring in multiple progressive waves of gene expression. Major enriched GO terms for each developmental stage (Stage 1 - 5) are labeled in gray boxes. The upper panel shows the distribution of cell density across pseudotime. More details on DE genes can be found in Table S2. **(E)** A GRN displaying key regulators during mesenchymal cell differentiation. The GRN consists of 36 TFs expressed dynamically across pseudotime. Arrows in orange indicate activation, while blue represents repression. Node size indicates the number of predicted connections. The stages (Stage 1 - 5) correspond to Figure 2D. GRN, gene regulatory network.

To further clarify the genetic coordination during the process of mesenchymal cell differentiation, we analyzed the gene regulatory network using the expression of transcription factors (TFs) (Figure 2E). Across pseudotime, 36 TFs were identified in the 5 stages of differentiation. We found that many well-known TFs involved in palatal formation in mesenchyme were enriched, including *Osr1*, *Osr2*, *Foxf2*, *Pax9*, *Dlx5*, *Barx1* and *Msx1* [6, 27]. Notably, we observed the enrichment of high-mobility group box family (*Hmgb1*, *Hmgb2*, and *Hmgb3*) in Stage 4, which were not previously reported to be involved in palatogenesis, suggesting that these HMGBs might represent novel TFs involved in palatal development (Figure 2E).

### Subpopulations and dynamic differentiation of epithelial cells

Epithelial cells covering the surface of the mesenchyme, though less abundant than the mesenchymal cells, also play a crucial role in palate development [5]. Since the isolated tissues for scRNA-seq were obtained from regions near the eyes and nose (Figure S1), eye-related epithelial cells (C13 and C19) and nasal epithelial cells (C22) were identified based on the expression of their respective markers [28–32]. During the final stage of palatogenesis when hard tissues of teeth were forming, and a small group of cells, C28, were found to express dental cell markers, including *Ambn*, *Amelx,* and *Dspp* [33, 34]. We also observed that C5 and C7 highly expressed *Sox2*, which is a marker of pluripotent progenitor cells [35], suggesting that these two subpopulations might represent epithelial progenitor cells. *Krt14* is a marker for epithelial basal cells [36], and we found that C27 exhibited the highest expression of *Krt14*, thus identifying it as this category of cell type. Based on the expression of *Krt6a* and *Krt6b*, markers for periderm differentiation [14, 37], we initially speculated that C6 might represent periderm cells. However, given that periderm is a single-cell layer of cells and the over-representation of C6 in the whole epithelium (24.7%), we re-annotated C6 as *Krt6*+ cells (Figure 1D and E, Figure S3, Figure S4; Table S1).

In comparison to mesenchymal cells, the developmental trajectory of epithelial cells appeared to be more linear along pseudotime (**Figure 3**A and B, Figure S8). The progression of epithelial cells initiated from epithelial progenitor cells (C5 and C7), passed through epithelial basal cells (C27), reached the *Krt6*+ cells (C6), and eventually culminated in the formation of dental cells (C28) (Figure 3A-C). The dynamic process of epithelial cells suggested that C6, which might contain periderm cells, was a relatively mature cluster, whereas the dental cell C28 might develop from epithelial cells during palatal development, in agreement with prior studies [37, 42]. Similar to the analysis of pseudotime-dependent expression of DE genes in mesenchyme, the earliest stage of epithelial cell differentiation was characterized by expression of genes functionally related to cell proliferation, such as chromatin organization and mitotic cell cycle. As pseudotime progressed, genes involved in epithelial cell differentiation, and signaling of Erk1/2 and tyrosine kinases became more active. During Stage 3, we found an enrichment of GO terms associated with protein synthesis, such as translation elongation and cytoplasmic ribosomal proteins, and observed a decrease in GO terms related to proliferation. Stage 4 was primarily featured by the formation of cornified envelope, accompanied by positive regulation of cell death. In the final stage of epithelial differentiation during palatogenesis, accessory organelles of the epithelia cells, such as cilium, began to form, and genes associated with dental cells were enriched (Figure 3D; Table S3).

**Figure 3.**
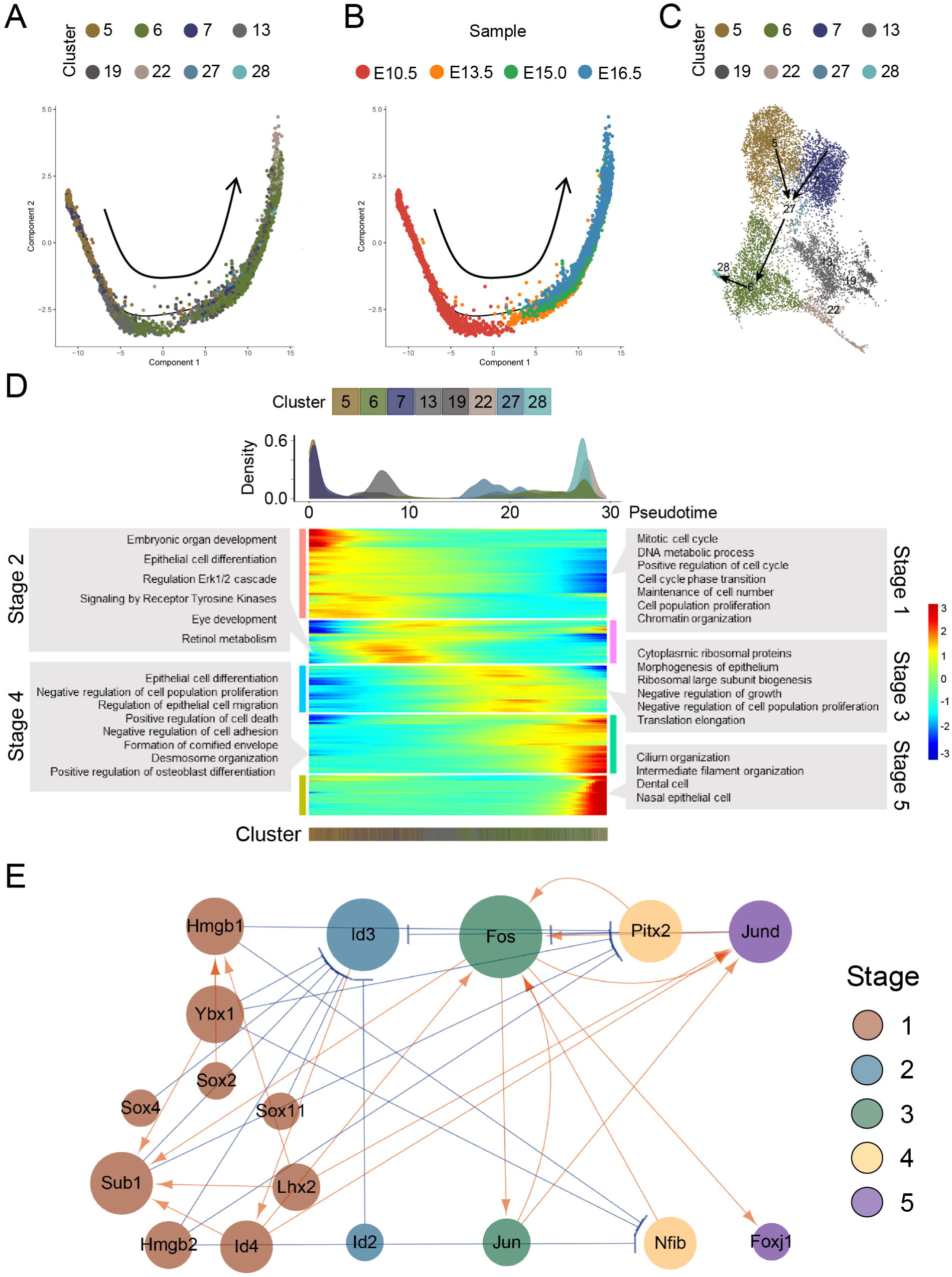
Characterization of dynamic differentiation of epithelial cells during palatogenesis in mice. **(A and B)** Pseudotime reconstruction of epithelial development reveals a linear ordering of cells. The cells are colored by **(A)** clusters and **(B)** time points. The arrow indicates the direction of developmental trajectories. **(C)** UMAP visualization of the epithelial trajectory colored by cell clusters. Arrows denote the trajectories of epithelial cells, initiating from epithelial progenitor cells, passing through epithelial basal cells, reaching the *Krt6*+ cells, and eventually forming dental cells. **(D)** Expression heatmap of 788 dynamically expressed genes ordered across pseudotime, showing epithelial differentiation occurring in multiple progressive waves of gene expression. Major enriched GO terms for each developmental stage (Stage 1 - 5) are labeled in gray boxes. The upper panel shows the distribution of cell density across pseudotime. More details on DE genes can be found in Table S3. **(E)** A GRN displaying key regulators during epithelial cell differentiation. The GRN consists of 17 TFs expressed dynamically across pseudotime. Arrows in orange indicate activation, while blue represents repression. Node size indicates the number of predicted connections. The stages (Stage 1 - 5) correspond to Figure 3D.

The gene regulatory network of TFs during the five stages of epithelial cell developmental trajectory showed that TF regulation was more complex during the early stages of differentiation, whereas it became simpler during later stages of development, suggesting that mature epithelial cells exhibited greater functional specificity than mesenchymal cells (Figure 3E). As the pseudotime progressed, it was observed that *Id3*, a marker for the activation of proliferation and suppression of epithelial-mesenchymal transition (EMT) [38], was negatively regulated by most TFs of Stage 1, indicating that cell proliferation began to be inhibited, and the EMT process was about to start. Furthermore, during Stage 3, we found that *Fos*, a marker of cornified cell formation [39], was enriched and potently activated *Foxj1*, a marker of cilia development [40], suggesting a potential relationship between keratinization and ciliogenesis [41] (Figure 3E).

### Four subpopulations of periderm cells give rise to two distinct trajectories of cell fates

We were intrigued by the apparent contradiction between the relative scarcity of periderm cells and the over-representation of C6 in the whole epithelium. To address this issue, we re-clustered C6 into 11 subclusters and identified C6.5, which showed specific expression of *Lypd3* and *Krt6a* [14, 42, 43], as the genuine subpopulation of periderm cells (**Figure 4**A, Figure S9). We found that there was a total of 224 cells in C6.5, ranging from E13.5 to E16.5. Notably, we did not observe any cells from the sample taken at E10.5, consistent with the finding of a previous report indicating that periderm differentiation during palatogenesis of mice begins after E10.5 [7]. We also conducted an IF assay with the marker gene *Krt6a* for periderm cells [37]. At E10.5, no KRT6A-positive cells were observed. However, from E13.5 to E16.5, we found KRT6A-positive cells covering the surface of the epithelial cells as a single layer, verifying that C6.5 was the subpopulation of periderm cells (Figure S5C).

**Figure 4.**
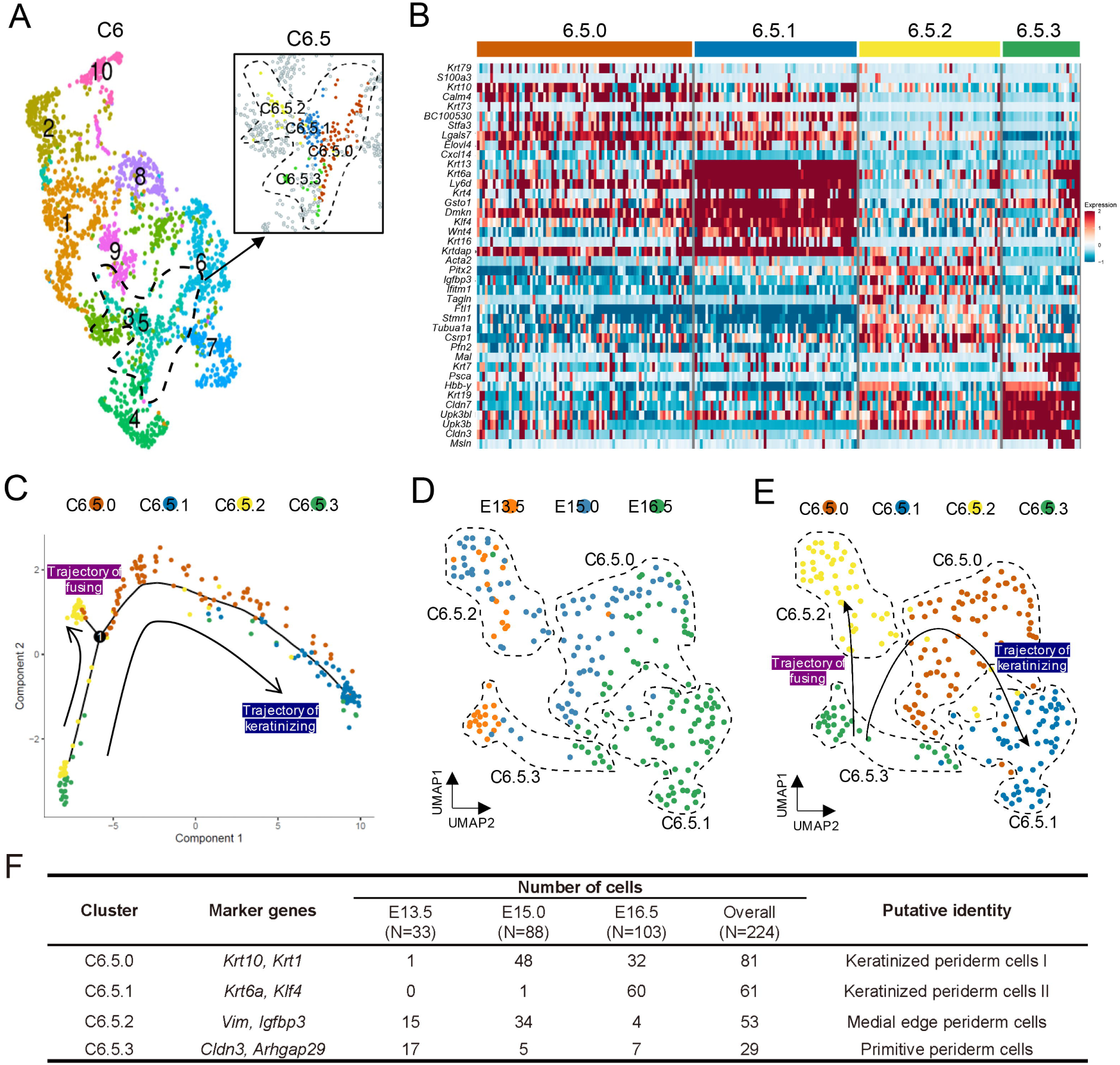
Identification of four subpopulations and two distinct trajectories of cell fate in periderm cells. **(A)** UMAP plot illustrating the re-clustering of the *Krt6+* population (C6), revealing C6.5 as the genuine subpopulation of periderm cells. Further subdivision of C6.5 into four subpopulations: C6.5.0, C6.5.1, C6.5.2, and C6.5.3 (shown in the zoom-in figure in the box). The dashed line indicates the periderm cell population C6.5. **(B)** Heatmap highlighting key marker genes utilized for inferring periderm cell subpopulations. The color scale represents the expression level. **(C)** Pseudotime reconstruction of periderm development reveals two distinct developmental trajectories originating from the same starting subpopulation C6.5.3. One trajectory leads to C6.5.2 (trajectory of fusing, highlighted in purple), while the other trajectory leads to C6.5.0 and C6.5.1 (trajectory of keratinizing, highlighted in blue). Arrows indicate the directions of the developmental trajectories. **(D and E)** UMAP visualization of the trajectory of periderm, colored by **(D)** time points and **(E)** cell clusters. Dashed lines indicate the four subpopulations of periderm cells. **(F)** Summary of subpopulations of periderm cells.

Previous studies have demonstrated that periderm covering palatal shelves exhibits unique gene expression patterns during palatogenesis [8]. In order to investigate the cellular and molecular transitions of palatal periderm, we further divided C6.5 into four subclusters (C6.5.0, C6.5.1, C6.5.2, and C6.5.3) with finer resolution (Figure 4A and B). Since no specific periderm marker genes at the single-cell resolution level were available, we identified these subpopulations based on their own gene sets and the composition of their developmental stages. We observed that C6.5.3, which mainly consisted of cells from E13.5, exhibited high expression of claudin-family protein coding genes (*Cldn3* and *Cldn7*) [14] and a well-known anti-adhesion protein coding gene (*Arhgap29*) [44]. Therefore, we referred that C6.5.3 represented the periderm cells covering the palatal shelves before they made contact with each other, annotating this population “primitive periderm cells”. We identified C6.5.2 as “medial edge periderm cells” due to their high expression of *Vim*, a gene known to be expressed in the medial edge epithelia (MEE) during the fusion of palatal shelves [45]. Finally, C6.5.0 and C6.5.1, which displayed high expression levels of various keratin coding genes, such as *Krt10*, *Krt1* and *Krt6a*, were identified as “keratinized periderm cells” [46]. We further annotated C6.5.0 as type I and C6.5.1 as type II, with type II being more mature than type I (Figure 4A, B, and F; Figure S10).

Through pseudotime analysis, it became apparent that these four subpopulations of periderm cells followed two distinct developmental trajectories, both originating from the same starting subpopulation (C6.5.3). One trajectory led to C6.5.2, which represented the structure of the MEE during palatogenesis, and was therefore identified as “trajectory of fusing”. The other trajectory led to the subpopulations of C6.5.0 and C6.5.1, which consisted of keratinized periderm cells, and was thus named “trajectory of keratinizing” (Figure 4C-F). To validate the existence of the four subpopulations and two developmental trajectories identified in our scRNA-seq analysis, we conducted IF assays using specific marker genes for each subpopulation. Specifically, KRT10 was expressed in C6.5.0 (keratinized periderm cells I), KLF4 in C6.5.1 (keratinized periderm cells II), IGFBP3 in C6.5.2 (medial edge periderm cells), and ARHGAP29 & CLDN3 in C6.5.3 (primitive periderm cells) (Figure S11-S14). The IF results revealed that the expression patterns of these marker genes were consistent with the scRNA-seq data, confirming the presence of the four subpopulations and supporting the two developmental trajectories proposed.

Next, the expression of DE genes across pseudotime was analyzed in these two distinct developmental trajectories separately (**Figure 5**A and B). In the trajectory of fusing, periderm cells initially showed an enrichment of genes related to the formation of intermediate filaments, followed by the upregulation of genes associated with tight junction formation. In the final phase of the fusing trajectory, we observed the appearance of GO terms related to apoptosis and ubiquitin-protein, suggesting that periderm cells in this trajectory might undergo protein breakdown and cell death (Figure 5A). Compared to the fusing trajectory, the keratinizing trajectory displayed a higher abundance of genes related to keratinization and formation of the cornified envelope in the final stage (Figure 5B), indicating that periderm cells in the keratinizing trajectory underwent cell death through the formation of a stratum corneum, consistent with previous studies [8].

**Figure 5.**
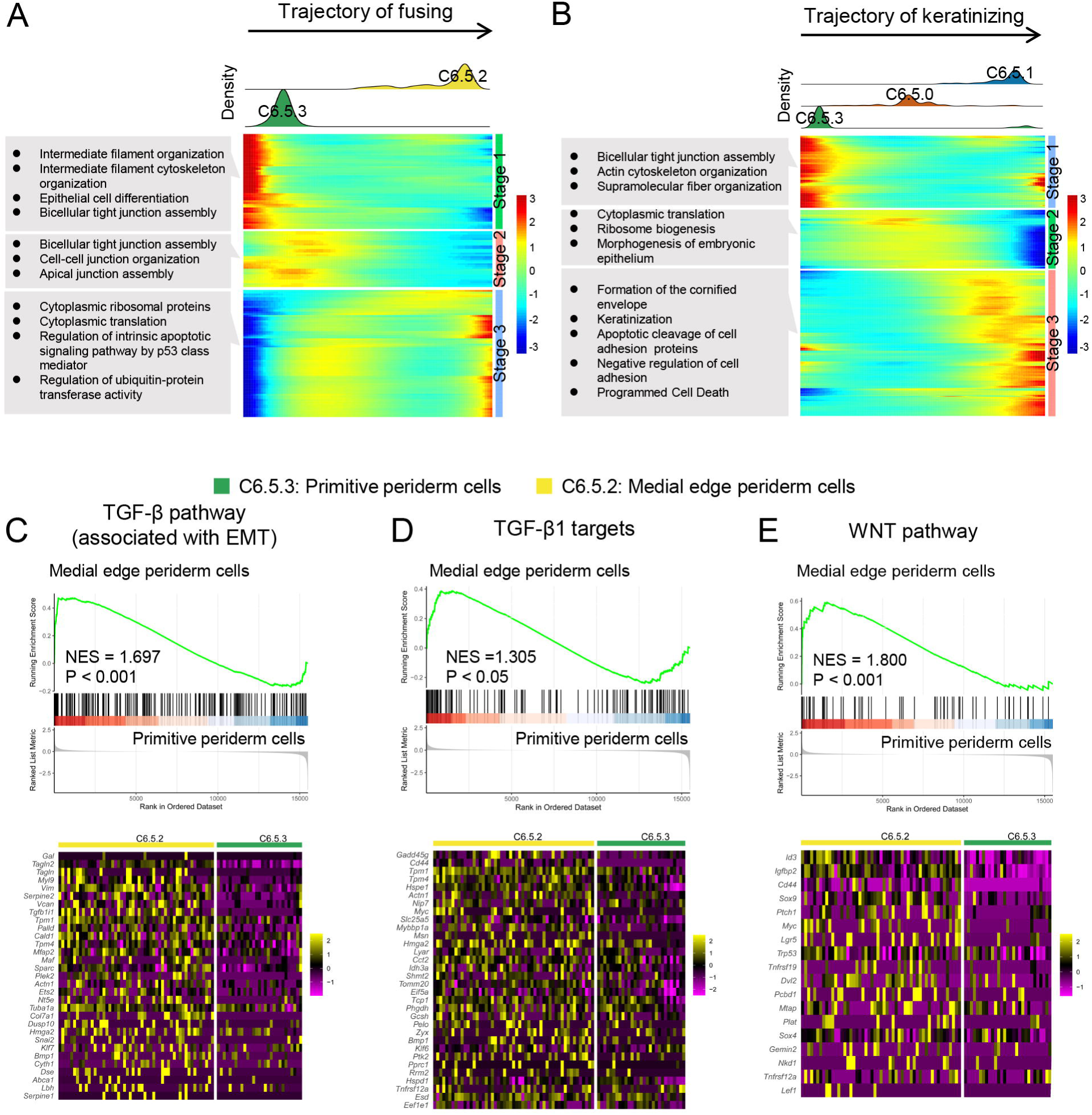
Characterization of dynamic differentiation of periderm cells during palatogenesis in mice. **(A and B)** Expression heatmap of 156 & 415 dynamically expressed genes ordered across pseudotime for **(A)** trajectory of fusing and **(B)** trajectory of keratinizing. Major enriched GO terms for each developmental stage (Stage 1 - 3) are labeled in gray boxes. The upper panel shows the distribution of cell density across pseudotime. **(C-E)** Signaling pathways enriched in medial edge periderm cells compared to primitive periderm cells by GSEA analysis. NES, normalized enrichment score. Lower panel, the corresponding heatmaps showing DE genes enriched in each pathway.

*Tgf-*β*3^-/-^*mice exhibit cleft palates caused by the failure of MEE degradation, highlighting the crucial role of TGF-β signaling in palatogenesis, especially in the fusion of palatal shelves [47]. Consistent with these findings, gene set enrichment analysis (GSEA) across pseudotime in the fusing trajectory revealed that TGF-β pathway components coding genes (associated with EMT and target genes) were upregulated in C6.5.2 (medial edge periderm cells) compared to C6.5.3 (Figure 5C and D). In addition, an enrichment of WNT pathway components coding genes was observed in C6.5.2 (Figure 5E), suggesting the important role of WNT signaling in the fusion.

In the analysis of the gene regulatory network of TFs in the trajectory of fusion, a total of 14 TFs were identified. Among them, *Pax9* was enriched in the first stage across pseudotime, which was consistent with its critical role in the regulation of the WNT pathway during palatogenesis [48]. Other TFs such as *Foxe1* and *Pitx2*, known to regulate TGF-β pathway [49, 50], were observed in later stages alongside the fusing trajectory (Figure S15A), corroborating the results obtained from GSEA. When examining the network of TFs in the keratinizing trajectory, we observed that *Fos* and *Klf4*, two important TFs required in the process of keratinization [16, 39], were enriched in the initial stage (Figure S15B), suggesting that a difference in the driving force between the keratinizing and fusing trajectories.

### *Claudins* and *Arhgap29* are involved in the non-stick function of the periderm prior to palatal shelves contact

Prior to the contact of opposing palatal shelves, the palatal periderm serves as a protective non-stick barrier, preventing pathological adhesion between the palatal shelves and other tissues [7]. Disruption of periderm, as observed in *Irf6*- or *Grhl3*-deficient mice, can cause abnormal oral adhesion and ultimately results in cleft palates [51, 52]. However, the detailed molecular mechanisms underlying the non-stick function of the periderm remain unclear. We observed high expression of various claudin-family coding genes, such as *Cldn3* and *Cldn7*, in primitive periderm cells (C6.5.3) (Figure 4B, Figure S10), which are main components of tight junctions. These tight junctions likely contribute to the exceptional tightness of the periderm-periderm connections in the monolayer of cells, effectively preventing exposure of adhesion molecules of the underlying epithelial cells. Moreover, *Arhgap29*, a molecule involved in anti-adhesion but necessary for periderm maintenance (like *Irf6* or *Grhl3*) [44], is specifically highly expressed in primitive periderm cells (Figure S10), suggesting that the periderm not only functions to insulate adhesion molecules of epithelia but also exhibits self-anti-adhesive properties.

We performed the co-expression analysis of *Arhgap29* and *Clnd3* in the C6.5.3 subpopulation and observed a concurrent expression of them (**Figure 6**A). We also investigated the expression of *Krt8* and *Clnd7*, two markers of anti-adhesion and tight junction [53, 54], and found similar co-expression pattern in primitive periderm cells (Figure 6B). Our scRNA-seq analysis was further confirmed by RNAscope *in situ* hybridization (ISH) on coronal sections of mouse craniofacial tissues at E13.5. We detected *Arhgap29* and *Cldn3* expression in the periderm region covering palatal shelves. Notably, *Arhgap29* was predominantly expressed on the apical and basal membrane of the periderm cells, while *Cldn3* was expressed at the lateral membrane of the periderm cells (**Figure 7**A and A’).

**Figure 6.**
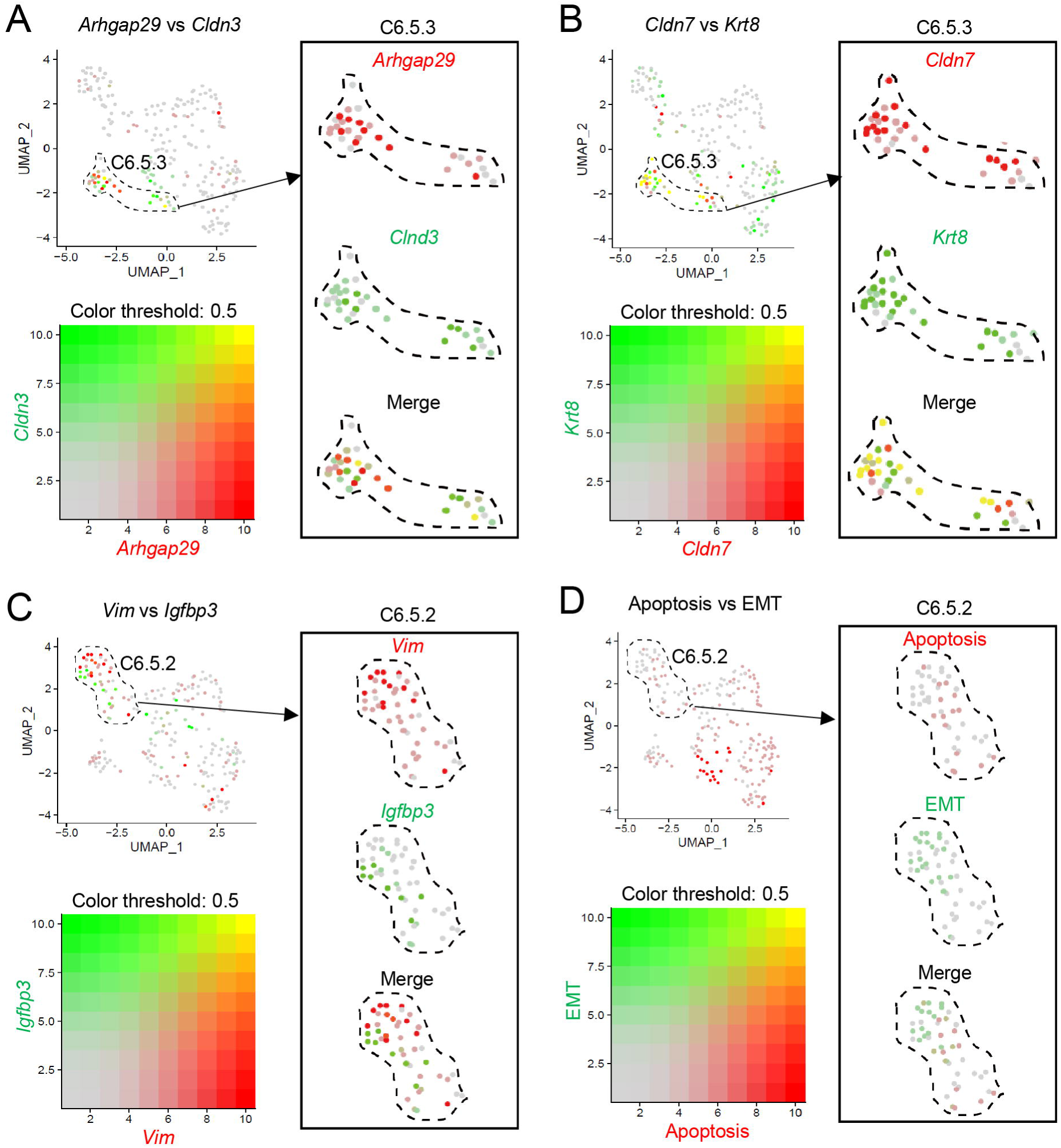
Co-expression patterns of marker genes for tight junctions and anti-adhesive molecules in primitive periderm cells C6.5.3, and mutual exclusivity expression patterns of EMT and apoptosis markers in medial edge periderm cells C6.5.2. **(A and B)** UMAP plots with overlaid expression of marker genes for tight junctions (*Cldn3* and *Cldn7*) and anti-adhesive molecules (*Arhgap29* and *Krt8*). Notably, the boxes magnify the co-expression of **(A)** *Arhgap29*&*Cldn3* and **(B)** *Krt8*&*Cldn7*. The dashed lines indicate the C6.5.3 subpopulation. **(C and D)** UMAP plots with overlaid expression of marker genes or gene sets for EMT and apoptosis. Notably, the boxes magnify the mutual exclusivity expression patterns of **(C)** *Vim*&*Igfbp3* and **(D)** EMT&apoptosis. The dashed lines indicate the C6.5.2 subpopulation. EMT, epithelial-mesenchymal transition.

**Figure 7.**
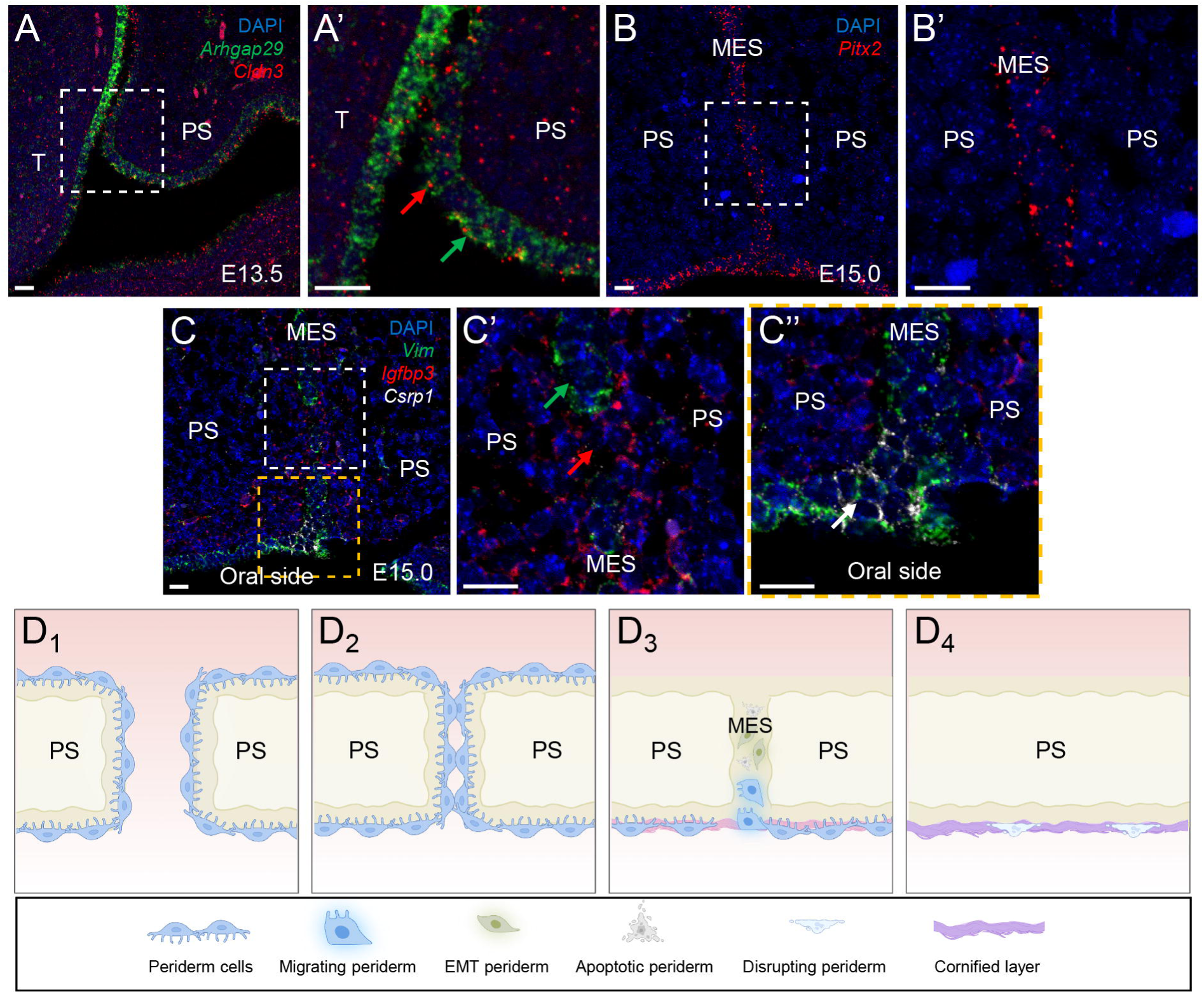
ISH validation of scRNA-seq and cellular transitions in peridermal cell fate during palatogenesis in mice. **(A and A’)** Coronal sections of E13.5 mouse embryos analyzed by RNAscope ISH, demonstrating expression of *Arhgap29* (green) and *Cldn3* (red) in the periderm region covering the palatal shelves. Notably, *Arhgap29* was primarily expressed on the apical and basal membrane of the periderm (green arrow in A’), while *Cldn3* was expressed at the lateral membrane of the periderm (red arrow in A’). **(B and B’)** Coronal sections of E15.0 mouse embryos analyzed by RNAscope ISH, showing *Pitx2* (red) expression alongside the MES of the fusing palatal shelves. **(C, C’ and C’’)** Coronal sections of E15.0 mouse embryos analyzed by RNAscope ISH, revealing the expression of *Vim* (green), *Igfbp3* (red), and *Csrp1* (white) in the MES of fusing palatal shelves. Notably, *Igfbp3* exhibited expression in the region where *Vim* did not fully link the entire MES (red and green arrows in C’), with rare co-localization of *Vim* and *Igfbp3*. *Csrp1* was predominantly expressed in the region of the MES abutting the oral side (white arrow in C’’). Dashed boxed areas are shown at higher magnification in A-C. Scale bars, 20 μm. T, tongue; PS, palatal shelf; MES, medial epithelial seam. **(D_1_)** Prior to the contact between opposing palatal shelves, the periderm functions as an anti-stick barrier by establishing tight junctions through claudins and expressing its own anti-adhesive molecules. **(D_2_)** Periderm cells at the medial edge start to guide adhesion. **(D_3_)** During the fusion phase, the periderm cells in MES disappear through a collective process of EMT, apoptosis and migration. **(D_4_)** The periderm cells covering the oral side (not in contact with the contralateral palatal shelf) undergo keratinization. PS, palatal shelf; MES, medial epithelial seam. Schematic representations of the cells are shown below D_1-4_. ISH, *in situ* hybridization.

Collectively, our results indicate that periderm acts as a protective non-stick barrier through two mechanisms: the formation of tight junctions via claudins to prevent the exposure of adhesion molecules of the underlying epithelial cells; the expression of anti-adhesive molecules such as *Arhgap29* to prevent adhesion.

### *Pitx2* mediates the adhesion of periderm during the contact of opposing palatal shelves

Periderm is believed to play a crucial role in guiding the adhesion of the medial edges of opposing palatal shelves [8, 10]. In the C6.5.2 subpopulation obtained mainly from E15.0 (the time point for palatal shelves adhesion), we observed specific expression of *Pitx2* (Figure 4B), which is known to be involved in adhesion functions [42]. This finding led us to speculate whether periderm cells undergo molecular transitions from anti-stick to guide-adhesion property at this developmental phase. To test this hypothesis, we employed RNAscope ISH and found that *Pitx2* was expressed alongside the medial epithelial seam (MES) of the fusing palatal shelf (Figure 7B and B’). Then we conducted *ex vivo* functional experiments using cultured palatal shelves. In the negative control group, the paired palatal shelves adhered successfully. However, when PITX2 expression was knocked down using adenovirus, the normal adhesion process was impaired (Figure S16). These results suggest that periderm functions as guiding adhesion of opposing palatal shelves at E15.0 during mouse palatogenesis and *Pitx2* is likely to mediate this process.

### Epithelial-mesenchymal transition, apoptosis and migration collectively contribute to the degeneration of periderm cells in the medial epithelial seam

The well-timed degeneration of periderm cells is critical for the complete fusion of palatal shelves, but there is still debate on whether and how this process is mediated by EMT, apoptosis, migration or other cellular mechanisms [8]. We found that the medial edge periderm subcluster expresses high levels of EMT marker *Vim* [45], apoptosis marker *Igfbp3* [55], and migration marker *Csrp1* [56, 57] (Figure 4B and F, Figure S10), suggesting that multiple cellular transitions collectively contribute to the degeneration of the periderm in MES. An additional question was whether these cellular processes occur in the same or different cells. Co-expression analysis revealed that *Vim* and *Igfbp3* were mainly highly expressed in C6.5.2. Notably, expression of these two genes showed mutual exclusivity within the same cell, suggesting that multiple cellular processes occurring in different cells beget to degeneration of MES (Figure 6C). We also conducted co-expression analysis of EMT-geneset and apoptosis-geneset, which yielded similar mutual exclusivity patterns of EMT and apoptosis (Figure 6D).

In validating the results and investigating the involvement of migration, RNAscope ISH was performed to detect the expression of *Vim*, *Igfbp3*, and *Csrp1* in the MES of fusing palatal shelves. *Vim* expression in the MES indicated that periderm was undergoing EMT (Figure 7C and green arrow in Figure 7C’). We also observed *Igfbp3* in the region where *Vim* was not linked to the whole MES (Figure 7C and red arrow in Figure 7C’). Notably, there was rare merge signaling of *Vim* and *Igfbp3* (Figure 7C and C’), corroborating the mutual exclusivity of EMT and apoptosis during the degeneration of periderm. Additionally, *Csrp1* was predominantly expressed in the region of MES abutting the oral side (Figure 7C and white arrow in Figure 7C’’), suggesting that the periderm degeneration mediated by migration mainly occurs more laterally along MES. In addition, three more marker genes relevant to this biological process were selected. STMN1 (an EMT marker [58]) and ANXA6 (an apoptosis marker [59]) were expressed within the MES region, whereas TAGLN (a migration marker [60]) was predominantly expressed in the region abutting the oral side (Figure S17).

Taken together, our results indicate that EMT, apoptosis and migration collectively contribute to the degeneration of periderm cells in MES.

## Discussion

Cleft palate is a common developmental structural defect that may occur in isolation or in combination with other anomalies, and it is challenging to be detected prenatally using ultrasound [4]. Therefore, investigating the expression patterns of genes during palatogenesis and unveiling their functions are critical to understand the etiology of this disorder and facilitate prenatal molecular diagnosis. Our recent bulk-RNA sequencing during mouse palatogenesis revealed the involvement of important biological processes such as cell adhesion and ossification, as well as key molecules, such as *Cdh1* and *Grhl3* [61]. However, due to the heterogeneity of different cell populations, bulk-RNA sequencing may not capture the precise molecular regulatory details, especially for rare populations like periderm cells.

In this study, we conduct scRNA-seq to create a comprehensive single-cell atlas of mouse developmental palates and make five main findings. First, we systematically depict the single-cell transcriptomics of mesenchymal and epithelial cells during palatogenesis in mice, including subpopulations and differentiation dynamics. Second, we identify four subclusters of palatal periderm cells and construct two differentiation trajectories. Third, we provide evidence that *Claudins* and *Arhgap29* are involved in the non-stick function of the periderm before the palatal shelves contact. Fourth, *Pitx2* mediates the adhesion of periderm during the contact of opposing palatal shelves. Finally, we demonstrate that EMT, apoptosis and migration collectively contribute to the degeneration of periderm cells in the medial epithelial seam.

The four subpopulations and two differentiation trajectories of periderm cells we identified suggest a model of periderm development during palatogenesis (Figure 7D_1_-D_4_). Before the opposing palatal shelves make contact, the periderm acts as anti-stick barrier by forming tight junctions *via* claudins and expressing its own anti-adhesive molecules (Figure 7D_1_). Upon contact of the opposing palatal shelves, the periderm covering the oral side (not in contact with the contralateral palatal shelf) undergoes keratinization (Figure 7D_3_ and D_4_), while the periderm cells within the medial edge transition from non-stick to guide-adhesion (Figure 7D_2_). At the fusion phase, the periderm cells in MES degenerate through a collective process of EMT, apoptosis and migration (Figure 7D_3_).

Based on this model, we notice that the key factor determining whether periderm undergoes adhesion-fusion or keratinization is the mode of contact between periderm cells. If periderm cells covering palatal shelves come into “face-to-face” contact, they undergo adhesion and then fuse together. However, if the periderm cells covering the oral side do not make “face-to-face” contact with other periderm cells, they undergo keratinization. Previous experiments using *in vitro* palatal shelves culture support this theory, as isolated palatal shelves can successfully fuse into a complete palate as long as they can come into close contact [62].

Another interesting observation was that although *Pitx2*, an adhesion marker, is expressed not only in MES but also widely in the oral epithelium (Figure S18A), it does not result in any pathological adhesion. Particularly, at the junction site between maxillary and mandibular tissue (white arrow in Figure S18B), the two layers of epithelial cells are merging into a single layer, indicating adhesion and fusion occur in this area. Based on these observations, we propose that close physical contact is the primary driving force of adhesion. When the opposing palatal shelves make contact, the force generated from underlying mesenchymal growth pushes the two palatal shelves to create close contact, and with the help of adhesion molecules, adhesion ultimately occurs. In the future, additional evidence from the isolated palatal shelves culture with varying levels of contact and discovery of mechanical signaling molecules will provide more insights into this hypothesis.

There has been controversy about the cellular mechanisms of degradation in MES during the fusion stage of palatogenesis [8]. Our study leveraged the advantages of scRNA-seq to analyze transcriptomic changes in individual cells and revealed that the degeneration of periderm in MES is mediated by EMT, apoptosis and migration in concert (Figure 6C and D, Figure 7C-C’’). Notably, co-expression analysis revealed that EMT and apoptosis exhibit mutual exclusivity within the same cell (Figure 6C and D, Figure 7C and C’), an intriguing observation not reported in previous studies.

Several recent studies have explored the biological processes involved in craniofacial development using scRNA-seq. Ozekin *et al.* [63] used scRNA-seq on E13.5 anterior palates, uncovering the corresponding epithelial and mesenchymal cell populations. Sun *et al.* [64] conducted scRNA-seq on maxillary prominences at E10.5, E11.5, E12.5, and E14.5, revealing a series of key transcription factors in the mesenchyme and the segregation of dental from palatal mesenchyme at E11.5. Jeremie *et al.* [65] performed single-cell RNA and ATAC sequencing on palate shelves at E13.5 and E15.5, and combined spatial RNA sequencing at E14.5 and E15.5, revealing spatiotemporally restricted expression patterns of key osteogenic marker genes through integrative analyses. Despite these advances, none of these studies included an analysis of periderm cells. Although Li *et al.* [14] examined single-cell transcriptomes in the fusion region of the upper lip and primary palate, they found that re-clustering of periderm cells did not reveal consistent subpopulations with obviously altered gene expression patterns, but only morphological changes. Therefore, none of these studies can explain the significant functional alterations in periderm cells.

When annotating cell clusters, in addition to the marker genes with known functional roles, we have also identified various genes that exhibit specific high expression but lack sufficient functional studies, such as *Upk3b*/*Upk3bl* in the C6.5.3 subpopulation (Figure 4B). These genes may also play important roles in palatogenesis, and exploring the functions of these genes *in vitro* or *in vivo*, along with evidence from clinical genetics, may yield novel insights into the etiology of cleft palate. Moreover, performing single-cell transcriptome analysis on the mouse model with a cleft, particularly focusing on novel candidate cleft genes without a clear molecular mechanism, and comparing these results to those from our current work on wild-type mice could provide more valuable findings, which require further investigation in future studies.

## Materials and methods

### Animal ethics

All animal studies were conducted in accordance with the guidelines and protocols approved by the Peking University Institutional Review Board (PUIRB-LA2018192), and the experiments were performed in strict accordance with the animal care.

### Microscopic isolation of tissues for scRNA-seq analysis

Wild type ICR mice were obtained from Beijing Charles River Laboratory Animal. Mouse embryos at E10.5, E13.5, E15.0 and E16.5 were selected according to our previous studies and others [6, 61]. Palates from five embryos at each time point were microscopically isolated, except for E10.5, for which the maxillary prominence and surrounding tissue were isolated. The isolation was performed in ice-cold PBS with 0.1% FBS (Catalog No. 10100147, Thermo Fisher Scientific, Grand Island, NY) solution for scRNA-seq analysis (Figure 1A, Figure S1).

### ScRNA-seq

The tissue was minced into small pieces using sterilized scissors and placed in 0.05% trypsin-EDTA solution (Catalog No. 25300054, Thermo Fisher Scientific), followed by gentle dispersion with a pipette. Then the tissue was dissociated at 37[, and Countess II (Catalog No. AMQAX1000, Thermo Fisher Scientific) was used to monitor the dispersion situation and vitality of cells to adjust the dissociation time. Once the tissue was dissociated into single cells with high viability (Figure S2A), the dissociation solution was quenched by adding complete medium containing FBS. The resulting single cell suspension was obtained by filtering the suspension through a 40-µm-porosity strainer (Catalog No. 352340, Falcon, Durham, NC), followed by centrifugation and resuspension in PBS containing 0.1% FBS.

The scRNA-seq libraries were prepared using the 10x Genomics Chromium Single Cell 3’ Reagent Kit v2 (Catalog No. 120264,10x Genomics, Pleasanton, CA), capturing cells at approximately 25% efficiency. After reverse transcription-polymerase chain reaction (RT-PCR), emulsions were broken and barcoded-cDNA was purified with dynabeads, followed by PCR amplification. To construct the 3’ gene expression library, the amplified cDNA was fragmented and end-repaired, and the DNA was double-size selected using SPRIselect beads. ScRNA-seq library sequencing was performed on the NovaSeq 6000 platform (Illumina, San Diego, CA) (Figure 1A, Figure S1).

### Dimensionality reduction and UMAP visualization

The sequencing reads were aligned to the Mus GRCm38 genome using CellRanger analysis pipeline v5.0. The Seurat R package (v4.1.0) was utilized for dimensionality reduction analysis [66]. The identification of variable genes in each cell was performed with default parameters, followed by principal component analysis (PCA) using the variable genes. To reduce dimensions and project different cell populations to two dimensions, uniform manifold approximation and projection (UMAP) was applied with a resolution value of 1.2.

### Identification of differentially expressed genes and cluster-specific marker genes

The FindAllMarkers function in the Seurat package (v4.1.0) [66] was used to identify the cluster-specific marker genes using normalized gene expression values of a given cluster to the average expression values in all other clusters, and FindMarkers function was used to identify differentially expressed (DE) genes between two clusters. The cutoff of average expression difference ≥ 0.25 natural Log and q < 0.01 was used.

### Analysis of single-cell developmental trajectory

The Monocle2 R package (v2.8.0) was utilized to perform pseudotime trajectory analysis using cluster marker genes identified in Seurat package (v4.1.0), as previously described [67]. Samples from different time points were used to determine the direction of the pseudotime trajectory. We employed the ‘plot_pseudotime_heatmap’ function with the default parameters to cluster the genes dynamically expressed along the pseudotime.

### GO analysis and gene set enrichment analysis

To characterize the various gene sets, gene ontology (GO) term analysis was performed using Metascape (https://metascape.org/) with default parameters [68]. In addition, gene set enrichment analysis (GSEA) was performed using the GSEA function in the R package ClusterProfiler version 4.2.2 [69].

### Analysis of gene regulatory network

The Monocle2 R package (v2.8.0) was utilized to normalize the expression levels of transcription factors (TFs) through the genesmoothcurve function, as previously described [67]. To infer reliable correlations, SCODE was employed to infer gene regulatory networks from dynamic TFs with a parameter z set to 4, and averaging 50x results [70]. Finally, we visualized the results using Cytoscape [71].

### RNAscope ISH

Mouse embryos were fixed in 4% PFA and then processed as coronal paraffin sections with a thickness of 4 μm. RNAscope^TM^ probes were designed according to the target genes and purchased from ACDBio. Hybridization and staining were carried out using the RNAscope Multiplex Fluorescent Reagent Kit v2 (Catalog No. 323100, ACDBio, Newark, CA) following the manufacturer’s protocol. DAPI was used to stain the nuclear nucleus. The stained sections were imaged using an Akoya PhenoImager HT imaging system and an Olympus FV3000 confocal laser scanning microscope.

### Immunofluorescence and immunohistochemistry assay

Coronal paraffin sections of mouse embryo heads underwent antigen retrieval and blocking. Then the sections were incubated overnight at 4°C with primary antibodies. Details of the primary antibodies for the marker genes are listed in Table S4. The following day, the sections were incubated with secondary antibodies and stained with DAPI for nuclear visualization. The immunohistochemistry (IHC) assay for PITX2 was conducted according to the manufacturer’s instructions (Catalog No. 67201-1-Ig, Proteintech, Rosemont, IL, 1:100). Images were captured using an automated whole-slide fluorescence scanner (Catalog No. Pannoramic DESK P-MIDI P250, 3DHISTECH, Budapest, Hungary).

### Palate culture *ex vivo* and adenovirus knock-down

To knock down *Pitx2*, we constructed an adenovirus containing shRNA targeting *Pitx2* (targeting sequence: 5’-CCGTACGTTTATAGGGACACATGTA-3’). A negative control adenovirus was used as previous reports [72]. We established an *ex vivo* palate culture system following previous methods [73, 74]. In brief, palatal shelves from E13.5 mouse embryos were dissected and placed in close contact on a 0.4 μm filter transwell within the lower chamber. The cultured palatal shelves were treated with DMEM/F12 medium (Catalog No. 11320033, Thermo Fisher Scientific) supplemented with 10% FBS (Catalog No. 10099158, Thermo Fisher Scientific) and 1% penicillin-streptomycin solution (Catalog No. 15140122, Thermo Fisher Scientific), with or without recombinant adenovirus. The cultured palates were incubated for 72 hours at 37°C, with the medium being changed every 24 hours. Then the cultured palates were fixed in 4% PFA, followed by paraffin embedding, and then underwent HE and IHC staining.

## Data availability

The raw sequence data reported in this paper have been deposited in the Genome Sequence Archive [75] in National Genomics Data Center [76], China National Center for Bioinformation / Beijing Institute of Genomics, Chinese Academy of Sciences (GSA: CRA019796) that are publicly accessible at https://ngdc.cncb.ac.cn/gsa.

## CRediT author statement

**Wenbin Huang**: Conceptualization, Validation, Data curation, Writing - original draft. **Zhenwei Qian**: Validation, Data curation, Writing - original draft. **Jieni Zhang:** Investigation, Validation, Data curation. **Yi Ding:** Methodology, Writing - review & editing. **Bin Wang:** Methodology, Data curation. **Jiuxiang Lin:** Conceptualization, Supervision, Writing - review & editing, Funding acquisition. **Xiannian Zhang:** Conceptualization, Supervision, Writing - review & editing, Funding acquisition. **Huaxiang Zhao:** Conceptualization, Data curation, Supervision, Writing - original draft, Writing - review & editing, Funding acquisition. **Feng Chen:** Conceptualization, Supervision, Writing - review & editing, Funding acquisition. All authors have read and approved the final manuscript.

## Acknowledgments

We thank Dr. Nan Jiang (Peking University) and Dr. Wenjie Zhong (Chongqing Medical University) for helping the analysis in this study. We thank Dr. Zhenzhen Fu (Nanchang University) for providing valuable information on the establishment of the *ex vivo* palate culture system. We thank Dr. Shiying Zhang, Dr. Mingzhao Li and Ms. Shujie Hou (Peking University) for their assistance in the capture of mouse embryos. This work is a part of a series of studies focusing on orofacial clefts under the supervision of Dr. Jiuxiang Lin. This work was supported by National Natural Science Foundation of China (No. 82170916, No. 82001030, No. 22104096, and No. 81870747), Shenzhen Fund for Guangdong Provincial High-level Clinical Key Specialties (No. SZGSP008), Shenzhen Clinical Research Center for Oral Diseases (No. 20210617170745001). Schematic illustrations were created with BioRender.com.

