## Supplementary material for "Single-cell atlas of developing mouse palates reveals cellular and molecular transitions in periderm cell fate": Figure S1-S18, Table S1, and Table S4

**Running title:** Single-cell landscape of mouse palatogenesis

Wenbin Huang[1, 7, 8, #], Zhenwei Qian[3, 4, #], Jieni Zhang[1, 9], Yi Ding[5], Bin Wang[2, 11], Jiuxiang Lin[1, 9, *], Xiannian Zhang[4, *], Huaxiang Zhao[2, 10, *], Feng Chen[6, 9, *]

### These authors contributed equally to this work.

**The supplemental materials include:**

✧ **Figure S1.** Single-cell RNA sequencing of 41,419 cells from the developing palate in mice across four critical developmental stages.

✧ **Figure S2.** Quality control and sequencing information.

✧ **Figure S3.** Heatmap highlighting relative expression levels of key marker genes across cell types and clusters.

✧ **Figure S4.** UMAP plots with overlaid expression of cluster-specific genes in mesenchymal cells, epithelial cells, and other cells.

✧ **Figure S5.** Immunofluorescence validation of mesenchymal, epithelial, and periderm cell populations identified by scRNA-seq.

✧ **Figure S6.** Distribution of cell clusters during palatogenesis in mice.

✧ **Figure S7.** Pseudotime reconstruction of mesenchymal development, illustrated separately by samples collected at different time points. Refer to Figure 2B.

✧ **Figure S8.** Pseudotime reconstruction of epithelial development, illustrated separately by samples collected at different time points. Refer to Figure 3B.

✧ **Figure S9.** Sub-clustering and annotation of C6 (*Krt6*+ cells).

- ✧ **Figure S10.** Characterization of marker genes expression in subclusters of periderm cells.
- ✧ **Figure S11.** Immunofluorescence validation of keratinized periderm cells I (C6.5.0) identified by scRNA-seq.
- ✧ **Figure S12.** Immunofluorescence validation of keratinized periderm cells II (C6.5.1) identified by scRNA-seq.
- ✧ **Figure S13.** Immunofluorescence validation of medial edge periderm cells (C6.5.2) identified by scRNA-seq.
- ✧ **Figure S14.** Immunofluorescence validation of primitive periderm cells (C6.5.3) identified by scRNA-seq.
- ✧ **Figure S15.** Gene regulatory networks (GRN) displaying key regulators during periderm cell differentiation.
- ✧ **Figure S16.** PITX2 knockdown impairs adhesion of mouse palatal shelves *ex vivo*.
- ✧ **Figure S17.** Immunofluorescence validation of EMT, apoptosis, and migration contributing to the degeneration of periderm cells in the medial epithelial seam.
- ✧ **Figure S18.** RNAscope *in situ* hybridization (ISH) demonstrating the expression of *Pitx2* in the MES as well as widely in the oral epithelium.
- ✧ **Table S1.** Summary of cell clusters during mouse palatogenesis.
- ✧ **Table S2.** The list of 479 differentially expressed (DE) genes across mesenchymal pseudotime, related to Figure 2D. (Presented in a separate Excel file)
- ✧ **Table S3.** The list of 788 differentially expressed (DE) genes across epithelial pseudotime, related to Figure 3D. (Presented in a separate Excel file)
- ✧ **Table S4.** Antibodies used in immunofluorescence assay for marker genes.

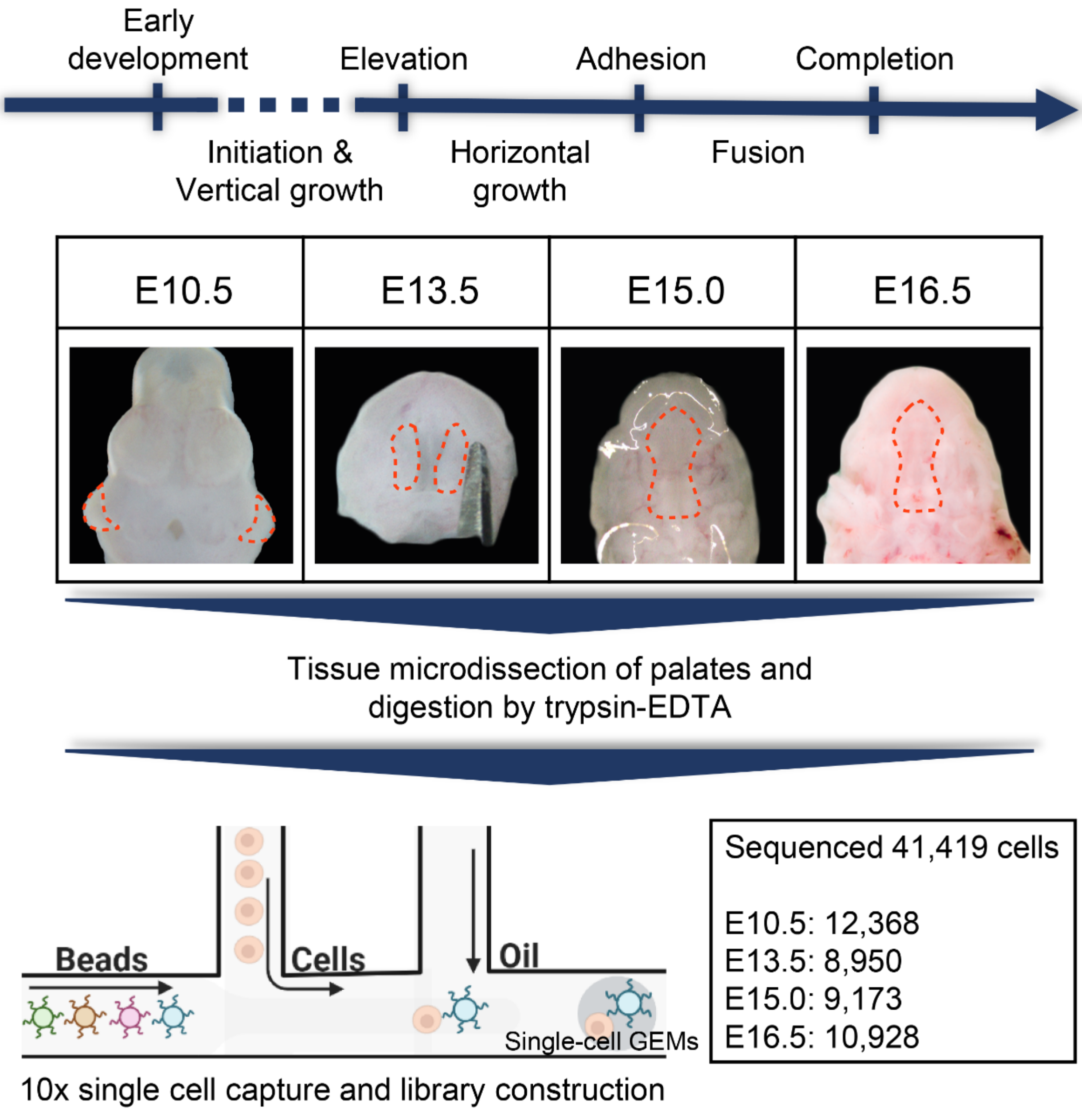


**Figure S1. Single-cell RNA sequencing of 41,419 cells from the developing palate in mice across four critical developmental stages.**

The stages include embryonic day 10.5 (E10.5, initial stage), E13.5 (vertical growth), E15.0 (fusion initiation) and E16.5 (completion). Palatal tissue at each stage was microscopically isolated, with the maxillary prominence and surrounding tissue isolated for E10.5. Subsequently, the isolated tissues were digested and subjected to scRNA-seq using the 10x Chromium system. The dashed circles denote the regions of isolated tissue in representative images of sampled embryos. Five embryos were microscopically isolated for analysis at each time point.

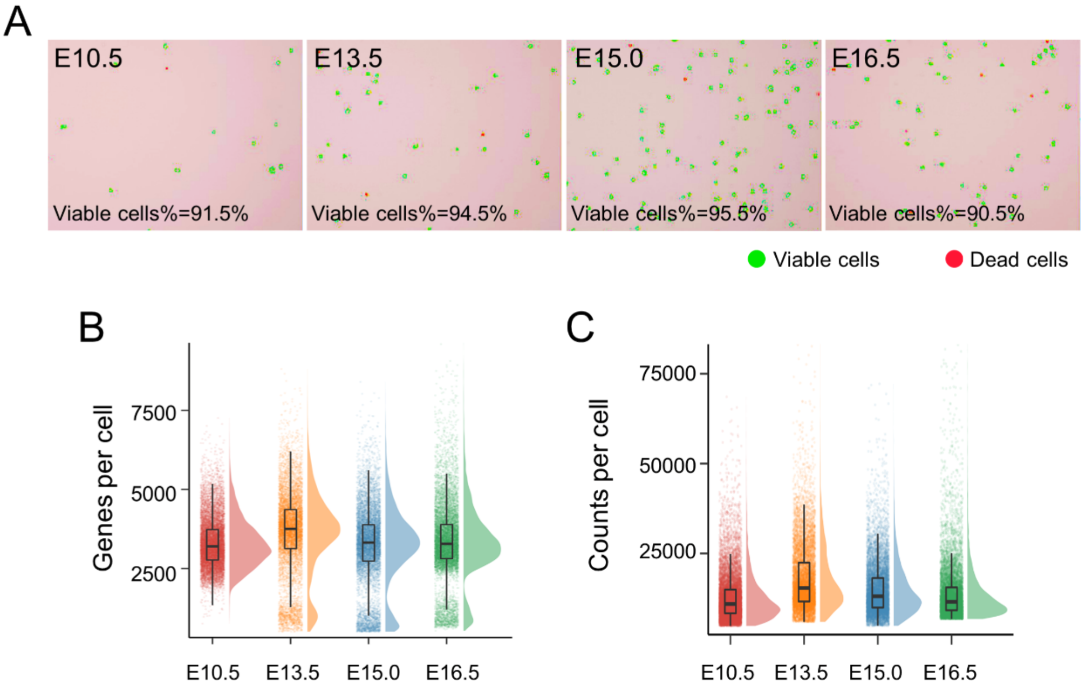


D

| Sample | Sequenced cells | Total reads | Reads Mapped to Genome | Sequencing saturation | Median Genes/cell | Median UMI/cell |
|---|---|---|---|---|---|---|
| E10.5 | 12,368 | 1,509,012,683 | 92.6% | 78.9% | 3,199 | 10,921 |
| E13.5 | 8,950 | 1,469,368,867 | 93.3% | 78.5% | 3,746 | 15,349 |
| E15.0 | 9,173 | 1,316,433,187 | 93.4% | 79.3% | 3,303 | 12,988 |
| E16.5 | 10,928 | 1,710,016,246 | 93.3% | 79.9% | 3,282 | 11,533 |

**Figure S2. Quality control and sequencing information.**

**(A)** Percentage of viable cells in samples at each time point following tissue digestion. **(B)** Number of detected genes per cell at each time point. **(C)** Number of detected UMI (unique molecular identifier) counts per cell at each time point. **(D)** Sequencing details for scRNA-seq of samples at four time points.

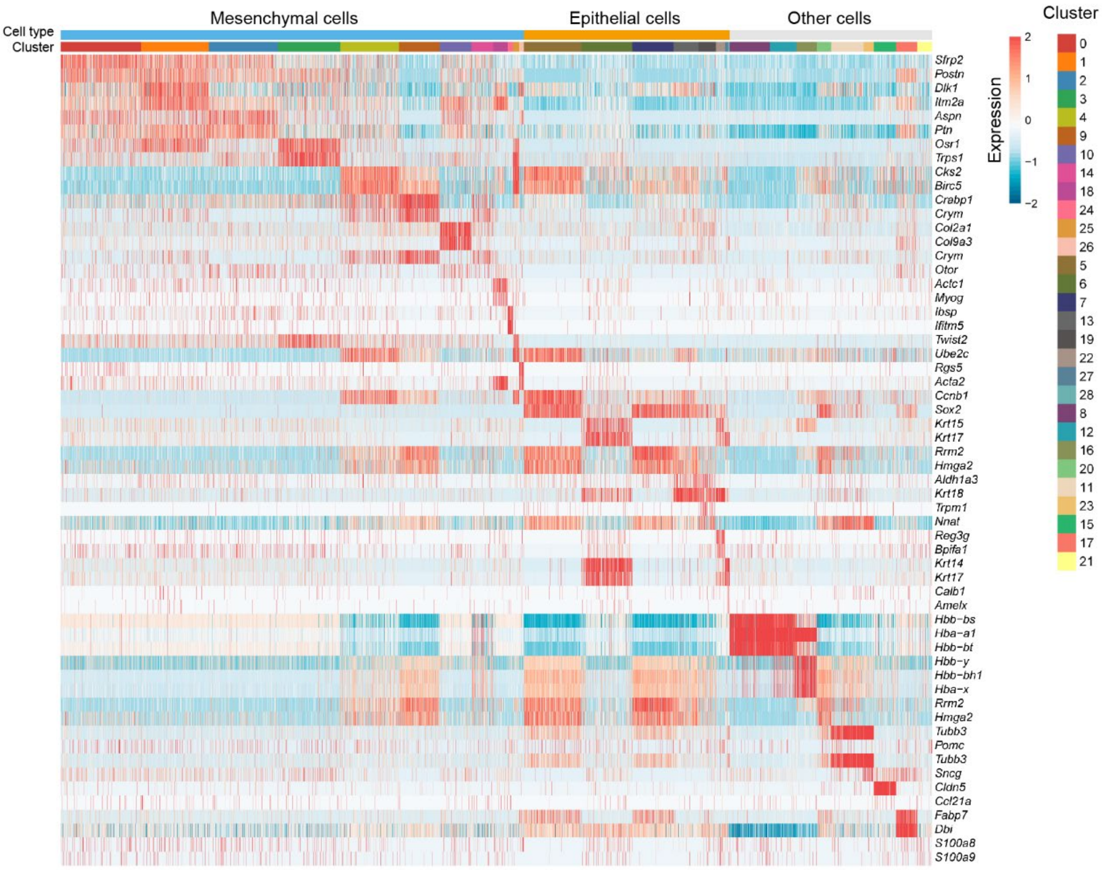


**Figure S3. Heatmap highlighting relative expression levels of key marker genes across cell types and clusters.**

The color scale represents the expression level. The upper bars indicate the major cell types, while the lower bars correspond to the cell clusters listed in Figure 1.

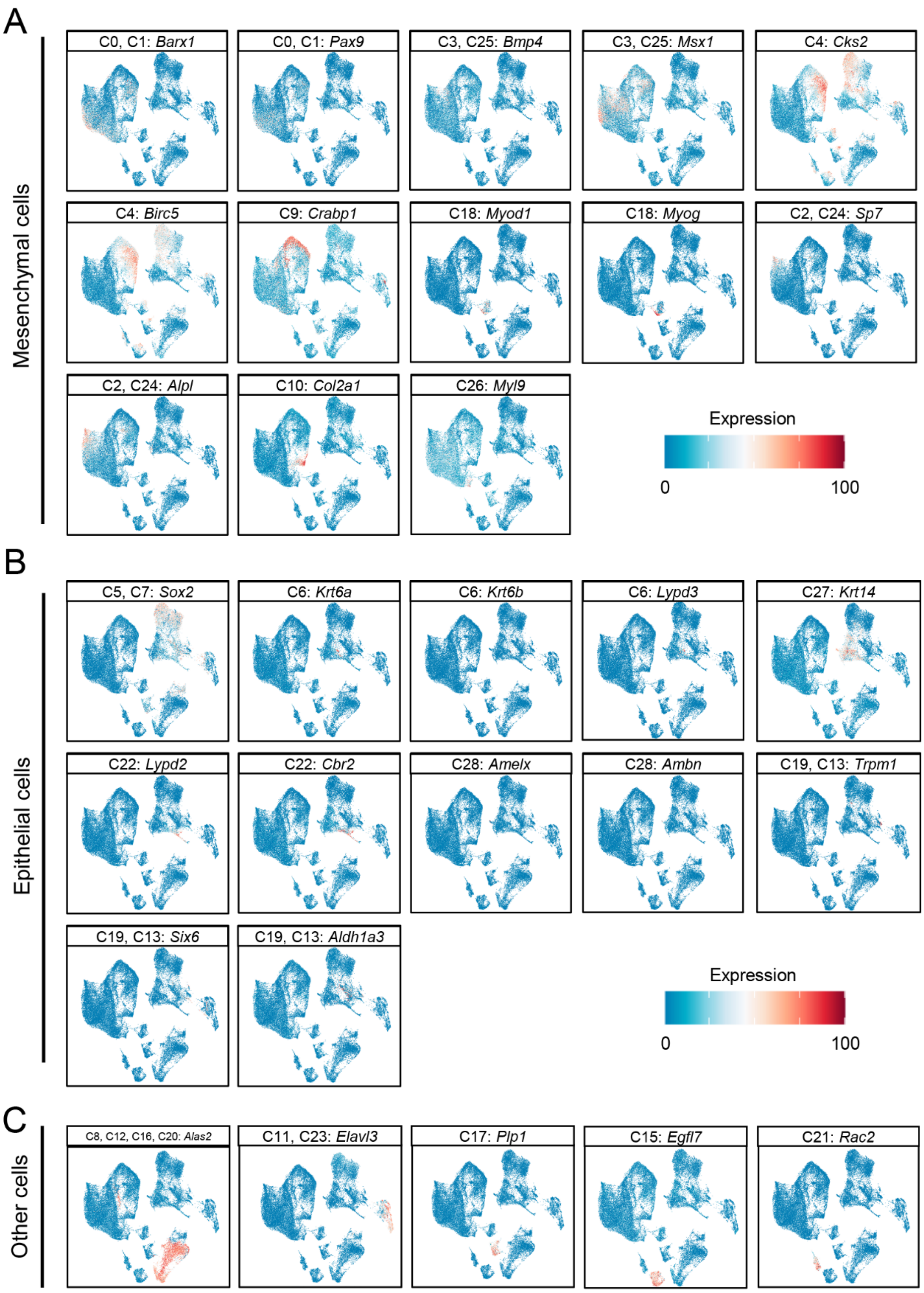


**Figure S4. UMAP plots with overlaid expression of cluster-specific genes in (A) mesenchymal cells, (B) epithelial cells, and (C) other cells.**

The color scale indicates the expression levels of the marker genes.

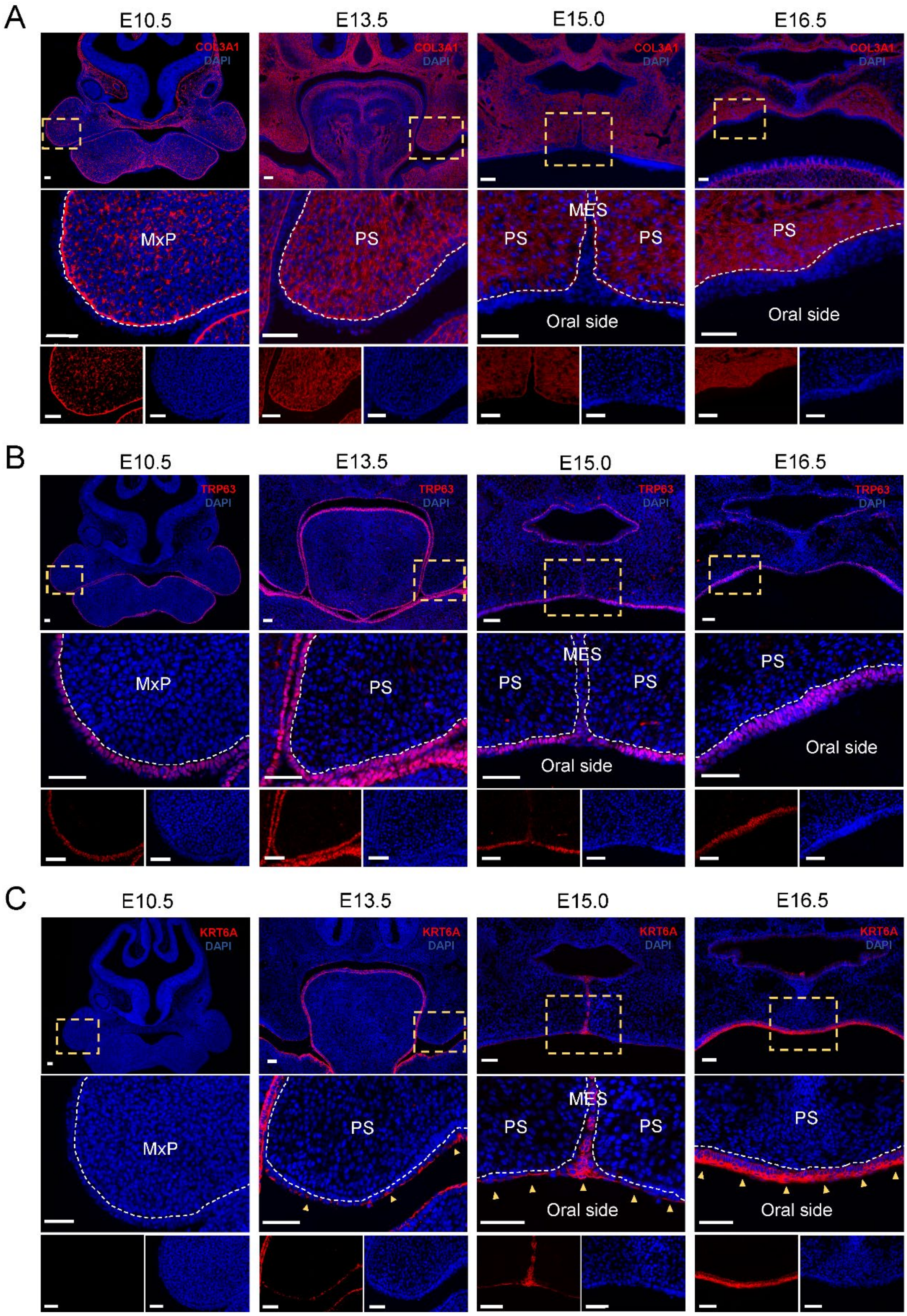


**Figure S5. Immunofluorescence validation of mesenchymal, epithelial, and periderm cell populations identified by scRNA-seq.**

Coronal sections of E10.5, E13.5, E15.0, and E16.5 mouse embryos were analyzed using immunofluorescence assays for **(A)** COL3A1 (marker gene for mesenchymal cell population), **(B)** TRP63 (marker gene for epithelial cell population), and **(C)** KRT6A (marker gene for periderm cell population). The results confirmed the presence of these cell populations. Notably, at E10.5, no KRT6A-positive cells were observed. However, from E13.5 to E16.5, there was a marked increase in KRT6A expression in periderm cells, consistent with our scRNA-seq analysis (refer to Figure S10). Dashed boxed areas are shown at higher magnification in the middle and bottom panels of (A), (B), and (C). White dashed lines indicate the boundary between the epithelium and mesenchyme, and yellow arrowheads indicate KRT6A-positive cells. Nuclei are stained with DAPI. Scale bars, 50 μm. PS, palatal shelf; MES, medial epithelial seam; MxP, maxillary prominence.

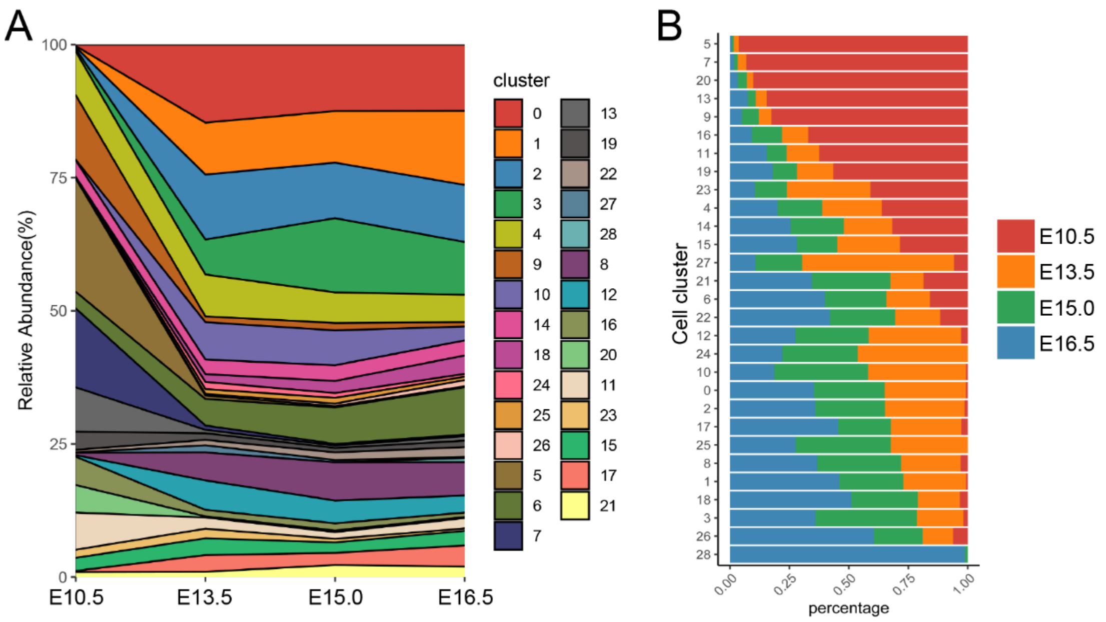


**Figure S6. Distribution of cell clusters during palatogenesis in mice.**

**(A)** Fraction of cell clusters at each time point, demonstrating an incremental cell-cluster complexity as the palate develops. **(B)** Bar plot illustrating the percentage of cells at each stage in each cell cluster, split by samples collected at different time points.

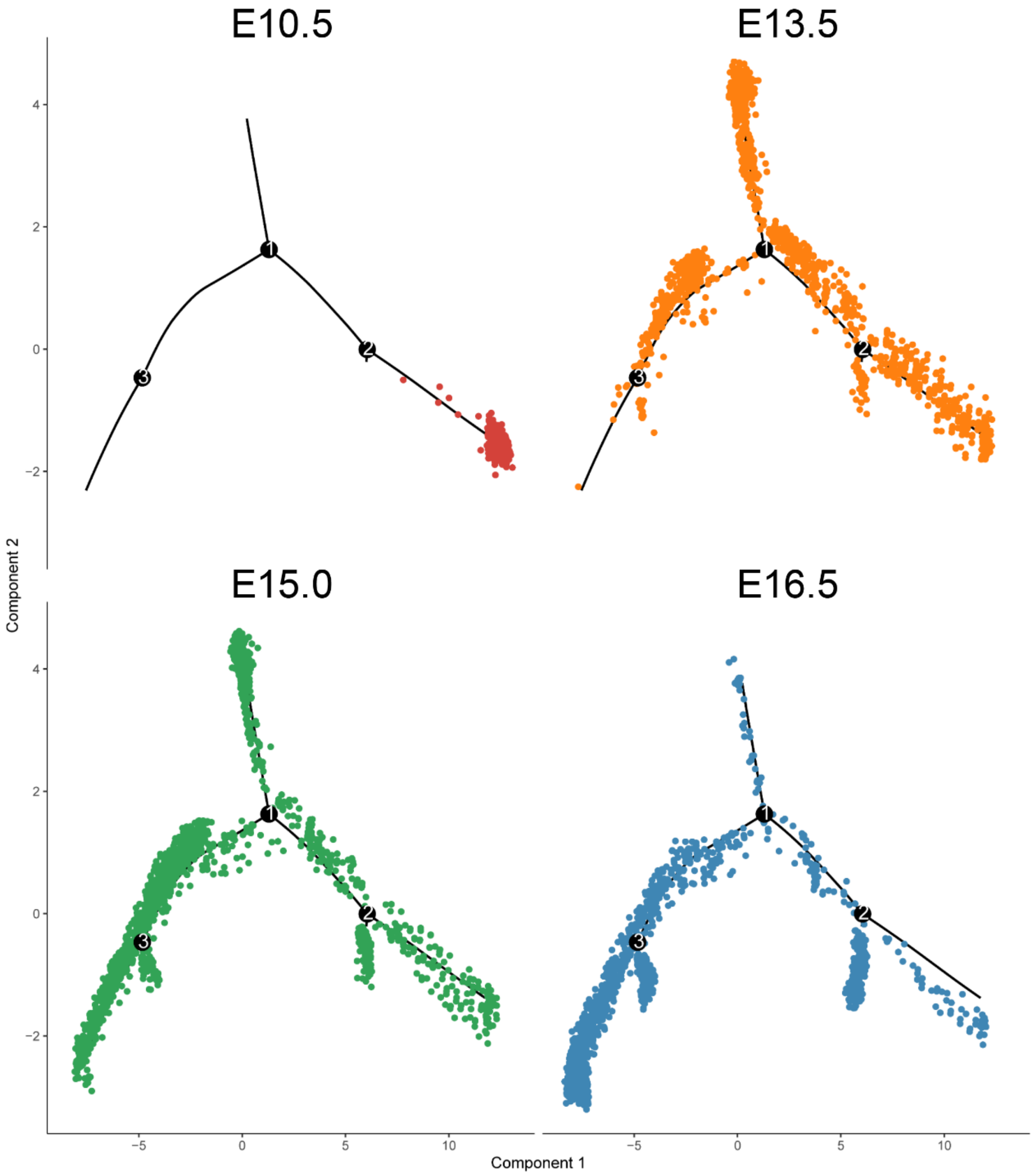


**Figure S7. Pseudotime reconstruction of mesenchymal development, illustrated separately by samples collected at different time points. Refer to Figure 2B.**

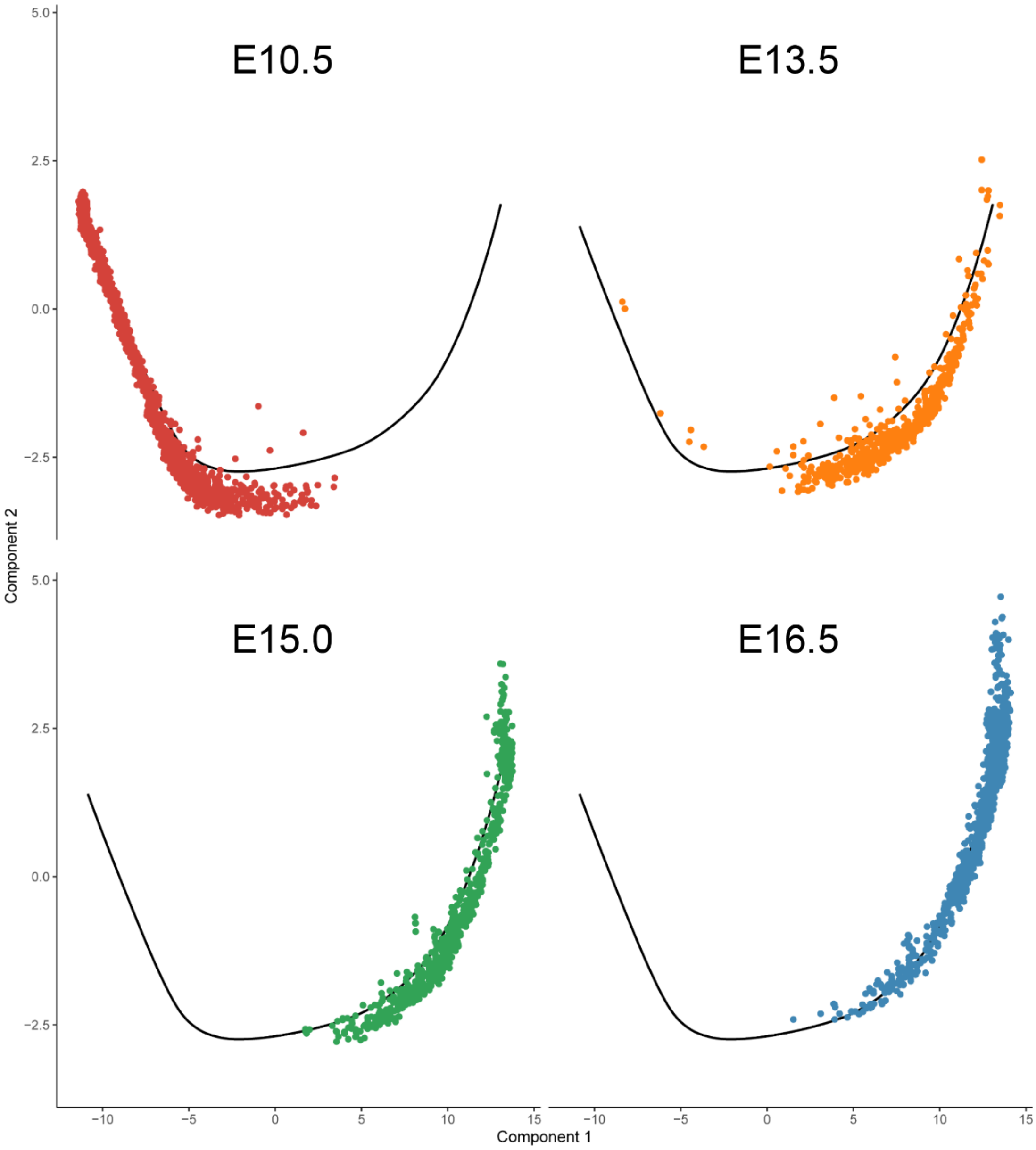


**Figure S8. Pseudotime reconstruction of epithelial development, illustrated separately by samples collected at different time points. Refer to Figure 3B.**

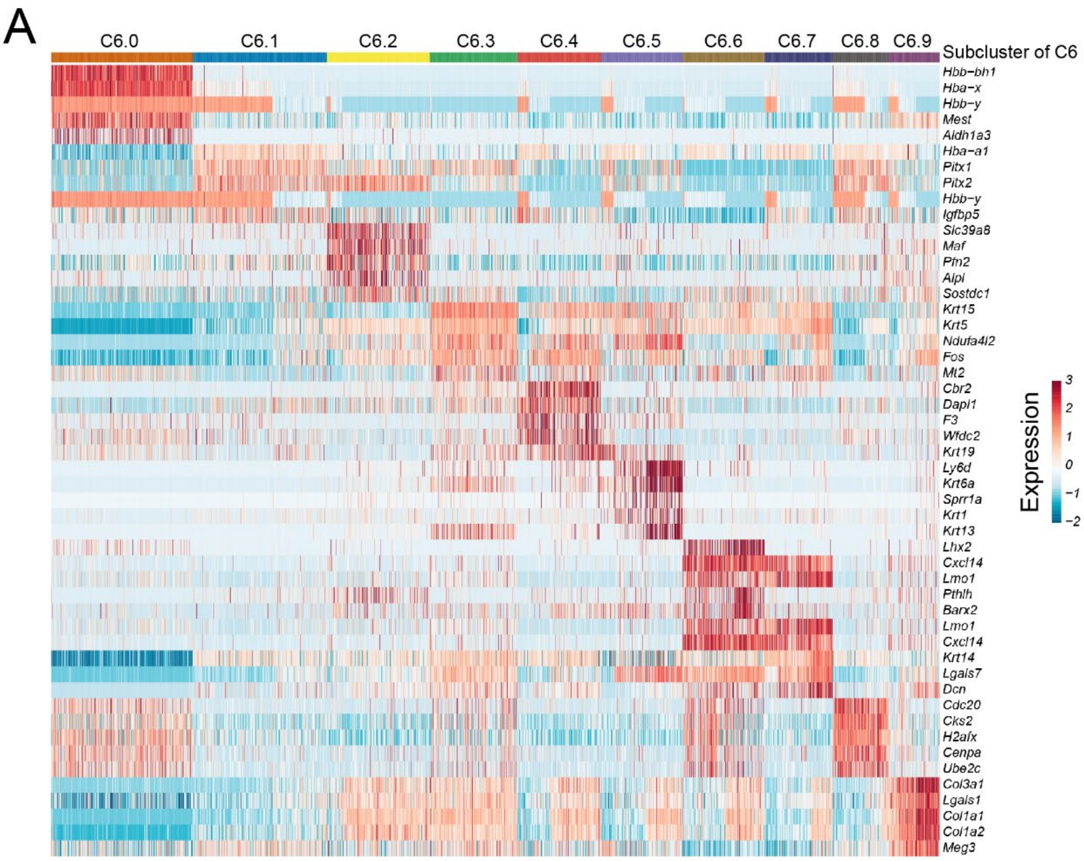


B

| Subcluster of C6 | Marker genes | Putative identity |
|---|---|---|
| C6.0 | *Hbb-bh1, Hba-x* | Red blood cells |
| C6.1 | *Hba-a1, Pitx1* | Ambiguous cell type |
| C6.2 | *Pthlh* | Dental follicle stem cells II |
| C6.3 | *Krt5* | Basal layer cells II-later stage |
| C6.4 | *Krt1, Krt10* | Spinous layer cells |
| C6.5 | *Krt6a, Lypd3* | Periderm cells |
| C6.6 | *Pthlh, Lhx2* | Dental follicle stem cells I |
| C6.7 | *Krt14, Krt5* | Basal layer cells I-early stage |
| C6.8 | *Cks2* | Early proliferating cells |
| C6.9 | *Lgals1, Postn, Bgn* | Ambiguous cell type |

**Figure S9. Sub-clustering and annotation of C6 (*Krt6*+ cells).**

**(A)** Heatmap highlighting key marker genes utilized for inferring periderm population in C6. The color scale represents the expression level. **(B)** Summary of subpopulations of C6.

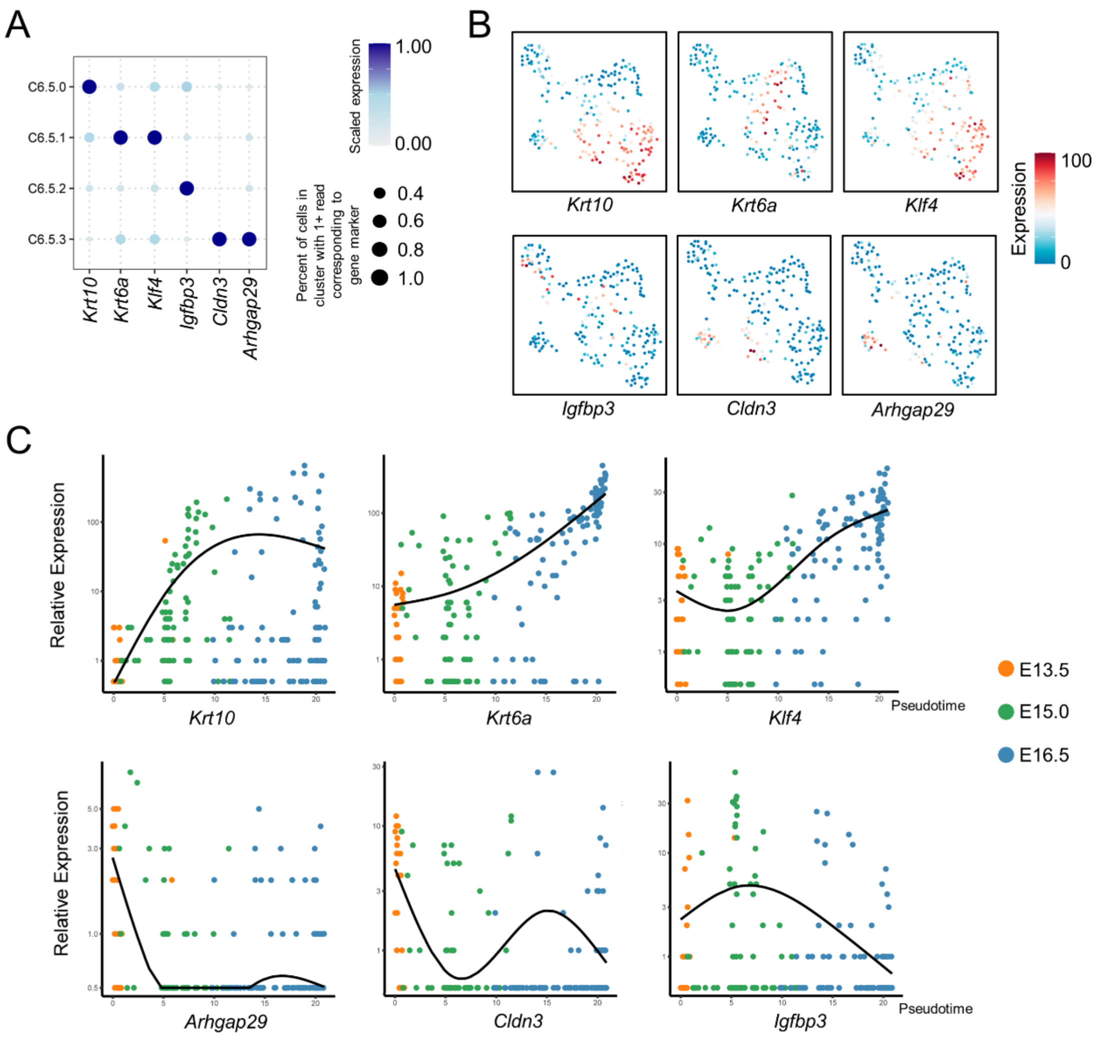


**Figure S10. Characterization of marker genes expression in subclusters of periderm cells.**

**(A)** Dot plot illustrating the relative expression of selected marker genes in each subcluster. The dot size represents the percentage of cells within a cell cluster where the relevant markers were detected. The color indicates the average expression level. **(B)** UMAP plots with overlaid expression of subcluster-specific genes, related to Figure 4D and E. The color scale indicates the expression level of the marker genes. **(C)** Kinetics plots displaying the relative expression of marker genes for periderm subpopulations across developmental pseudotime.

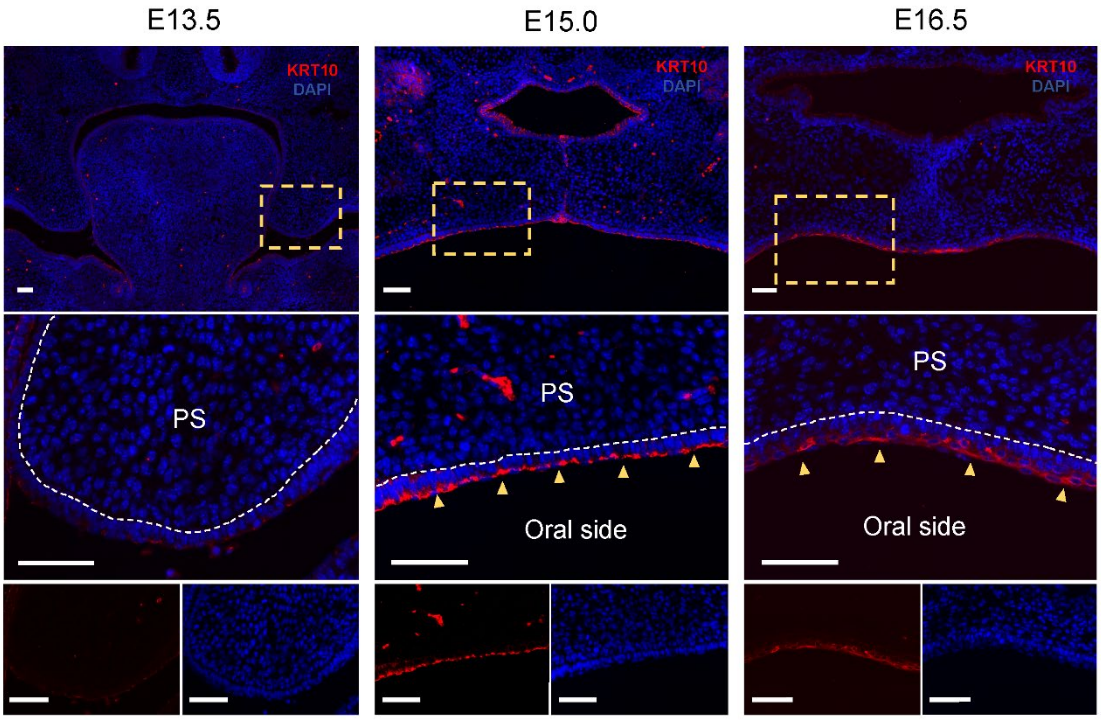


**Figure S11. Immunofluorescence validation of keratinized periderm cells I (C6.5.0) identified by scRNA-seq.**

Coronal sections of E13.5, E15.0, and E16.5 mouse embryos were analyzed using immunofluorescence assays for KRT10. At E13.5, no KRT10-positive cells were detected. However, at E15.0 and E16.5, numerous KRT10-positive cells were observed on the epithelial surface, with a slightly higher expression at E15.0 compared to E16.5. This expression pattern of KRT10 confirmed the identification of C6.5.0 as keratinized periderm cells I (refer to Figure 4 and Figure S10). Dashed boxed areas are shown at higher magnification in the middle and bottom panels. White dashed lines indicate the boundary between the epithelium and mesenchyme, and yellow arrowheads indicate KRT10-positive cells. Nuclei are stained with DAPI. Scale bars, 50 μm. PS, palatal shelf; MES, medial epithelial seam.

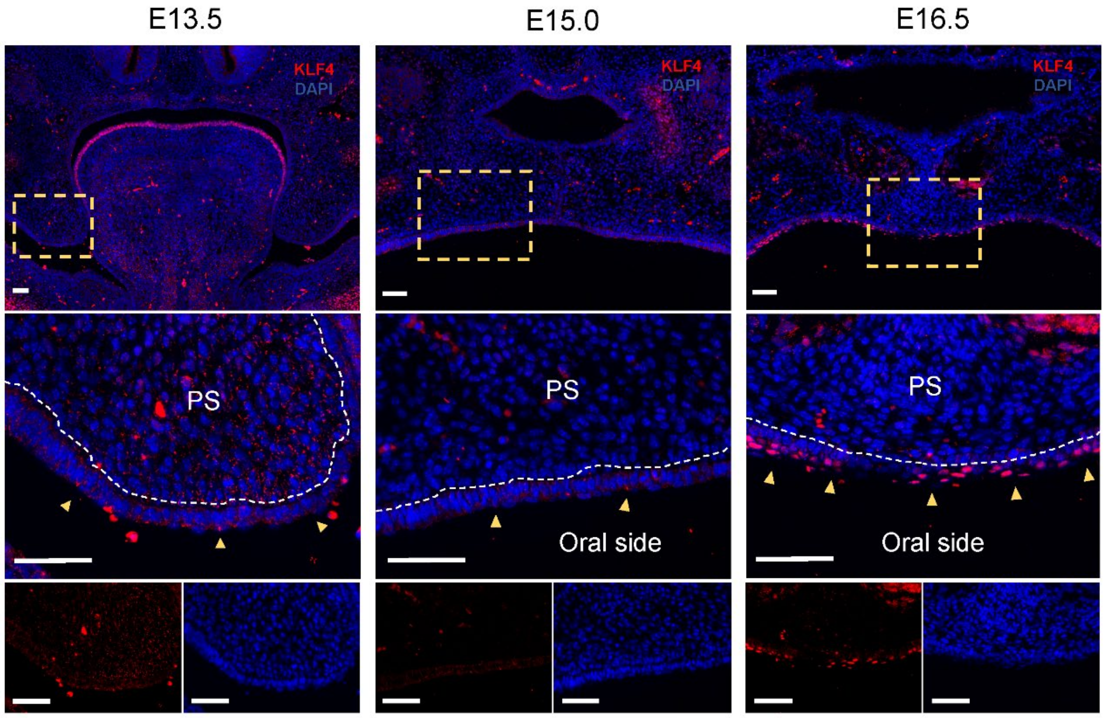


**Figure S12. Immunofluorescence validation of keratinized periderm cells II (C6.5.1) identified by scRNA-seq.**

Coronal sections of E13.5, E15.0, and E16.5 mouse embryos were analyzed using immunofluorescence assays for KLF4. At E13.5 and E15.0, only a few KLF4-positive cells were detected. However, at E16.5, there was a significant increase in KRT10-positive cells on the epithelial surface. The expression pattern of KLF4 was consistent with scRNA-seq analysis, confirming the identification of C6.5.1 as keratinized periderm cells II (refer to Figure 4 and Figure S10). Dashed boxed areas are shown at higher magnification in the middle and bottom panels. White dashed lines indicate the boundary between the epithelium and mesenchyme, and yellow arrowheads indicate KLF4-positive cells. Nuclei are stained with DAPI. Scale bars, 50 μm. PS, palatal shelf; MES, medial epithelial seam.

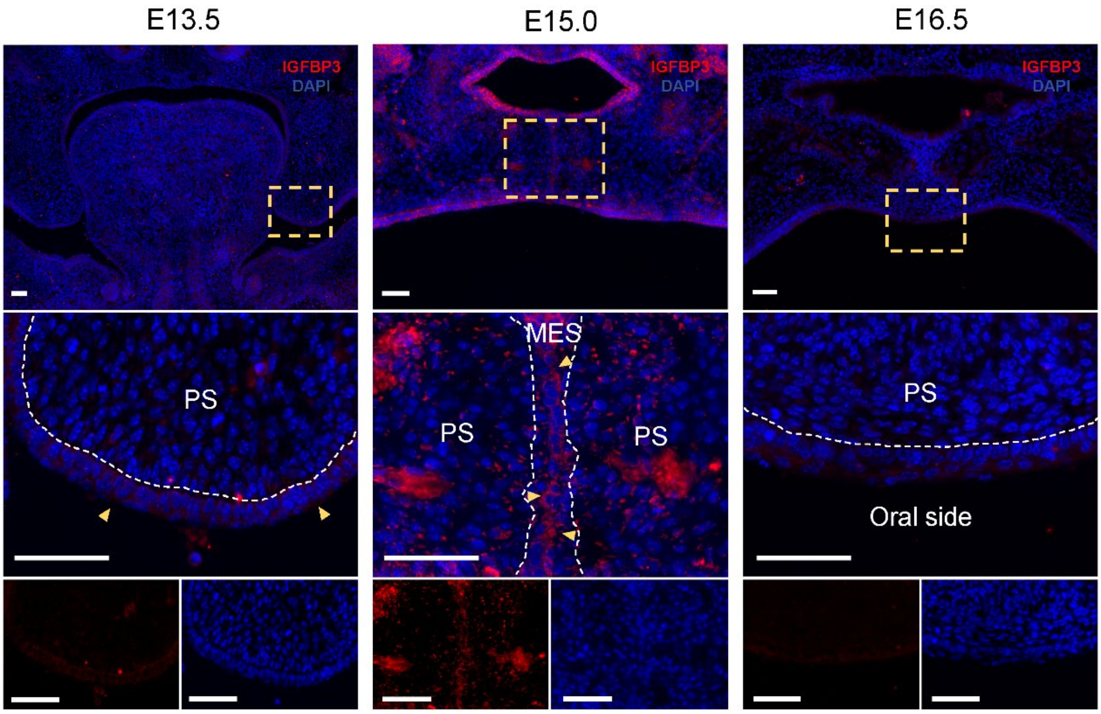


**Figure S13. Immunofluorescence validation of medial edge periderm cells (C6.5.2) identified by scRNA-seq.**

Coronal sections of E13.5, E15.0, and E16.5 mouse embryos were analyzed using immunofluorescence assays for IGFBP3. IGFBP3 was primarily expressed in the medial edge seam (MES) region at E15.0, while it was rarely detected in the epithelium region at both E13.5 and E16.5. These findings confirm the identification of C6.5.2 as medial edge periderm cells (refer to Figure 4 and Figure S10). Dashed boxed areas are shown at higher magnification in the middle and bottom panels. White dashed lines indicate the boundary between the epithelium and mesenchyme, and yellow arrowheads indicate IGFBP3-positive cells. Nuclei are stained with DAPI. Scale bars, 50 μm. PS, palatal shelf; MES, medial epithelial seam.

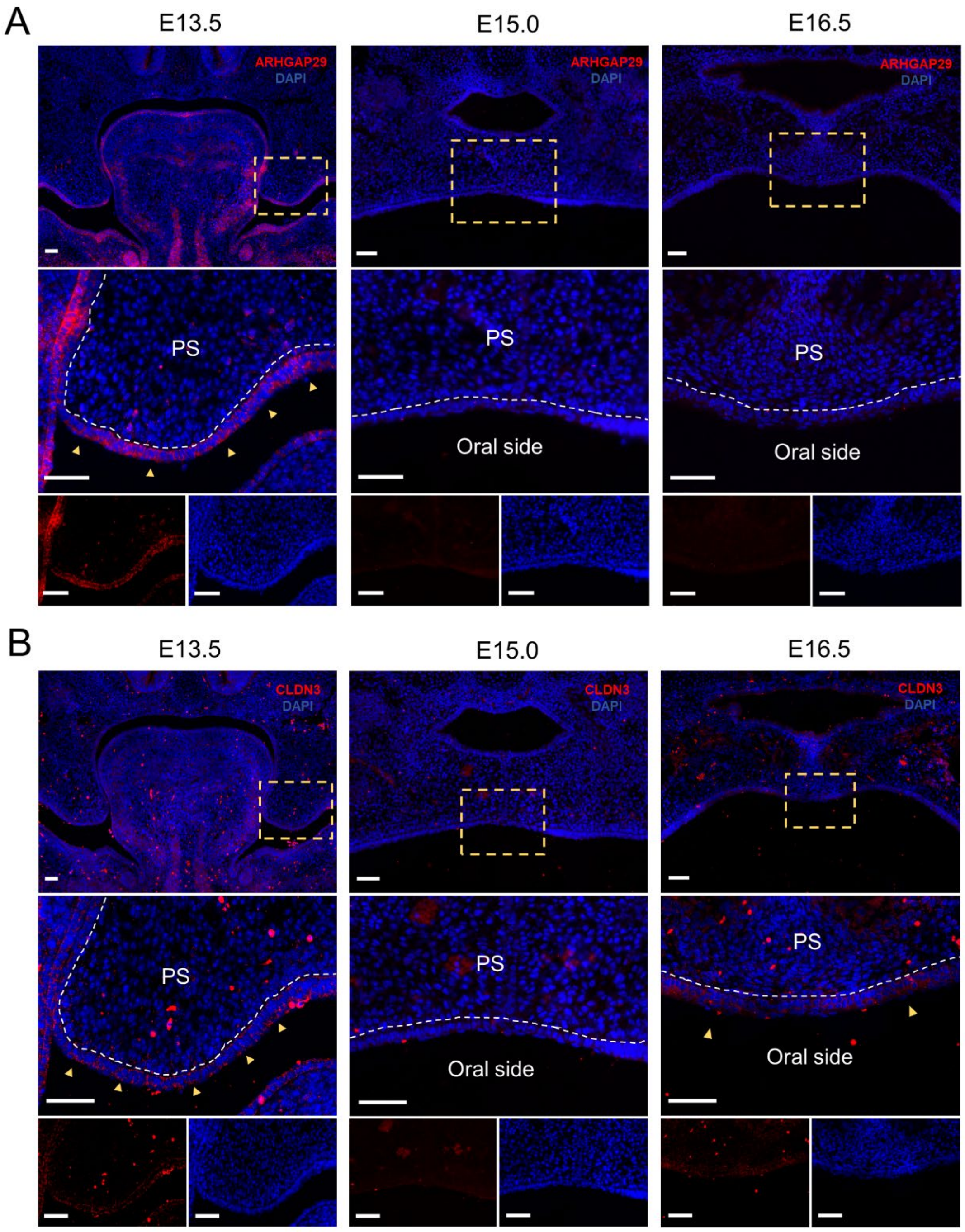


**Figure S14. Immunofluorescence validation of primitive periderm cells (C6.5.3) identified by scRNA-seq.**

Coronal sections of E13.5, E15.0, and E16.5 mouse embryos were analyzed using immunofluorescence assays for **(A)** ARHGAP29 and **(B)** CLDN3. At E13.5, both ARHGAP29- and CLDN3-positive cells were prominently observed on the epithelial surface of the palatal shelves. In contrast, at E15.0 and E16.5, few ARHGAP29- and CLDN3-positive cells were detected on the epithelial surface. The expression patterns of ARHGAP29 and CLDN3 confirmed the identification of C6.5.3 cells as primitive periderm cells (refer to Figure 4 and Figure S10). Dashed boxed areas are shown at higher magnification in the middle and bottom panels of (A) and (B). White dashed lines indicate the boundary between the epithelium and mesenchyme, and yellow arrowheads indicate ARHGAP29- and CLDN3-positive cells. Nuclei are stained with DAPI. Scale bars, 50 μm. PS, palatal shelf; MES, medial epithelial seam.

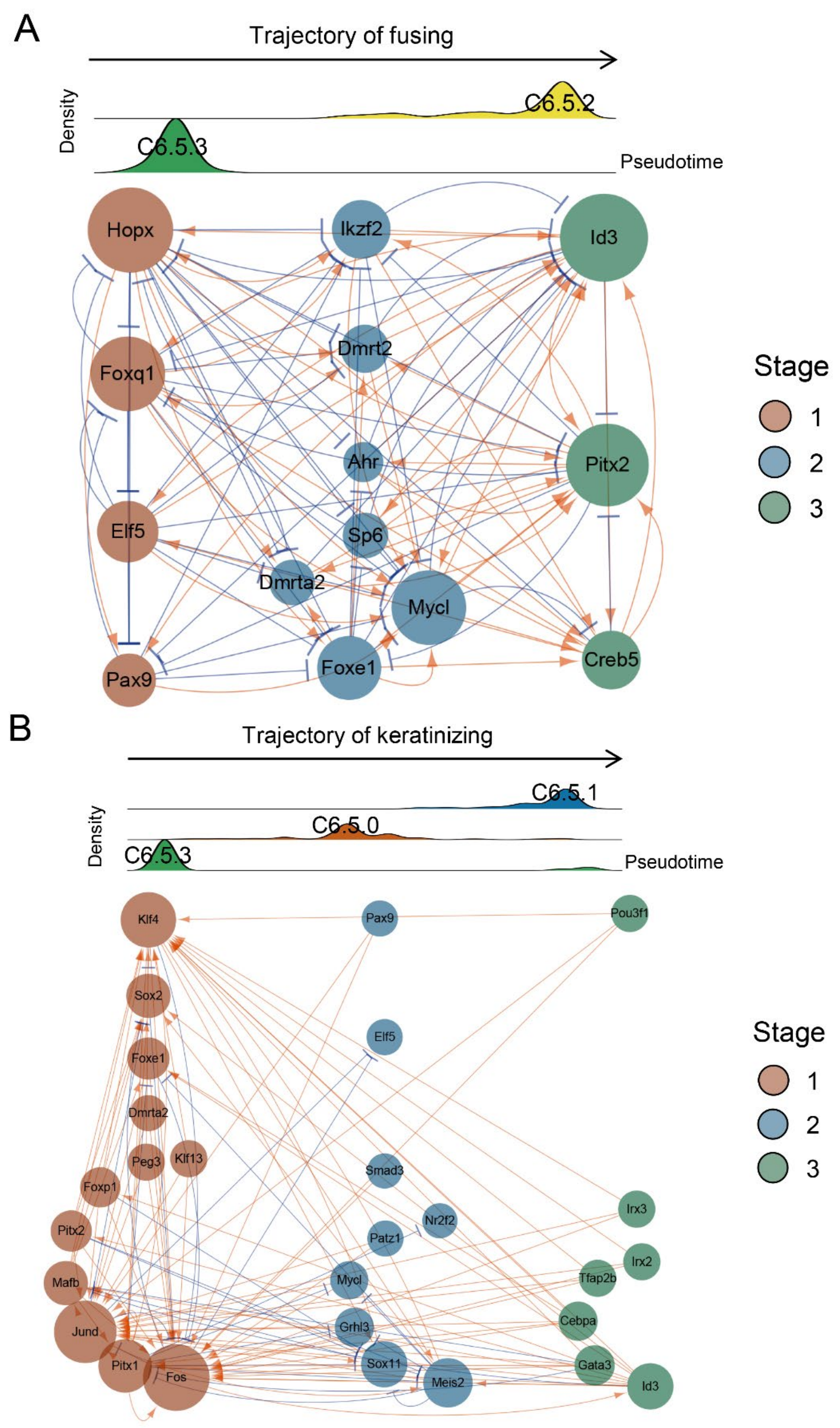


**Figure S15. Gene regulatory networks (GRN) displaying key regulators during periderm cell differentiation.**

**(A)** GRN for the trajectory of fusing and **(B)** GRN for the trajectory of keratinizing. The GRN consists of 14 & 28 TFs expressed dynamically across the pseudotime of the trajectory of fusing and the pseudotime of the trajectory of keratinizing, respectively. Arrows in orange indicate activation, while blue represents repression. Node size indicates the number of predicted connections. The stages (Stage 1 - 3) correspond to Figure 5A and B.

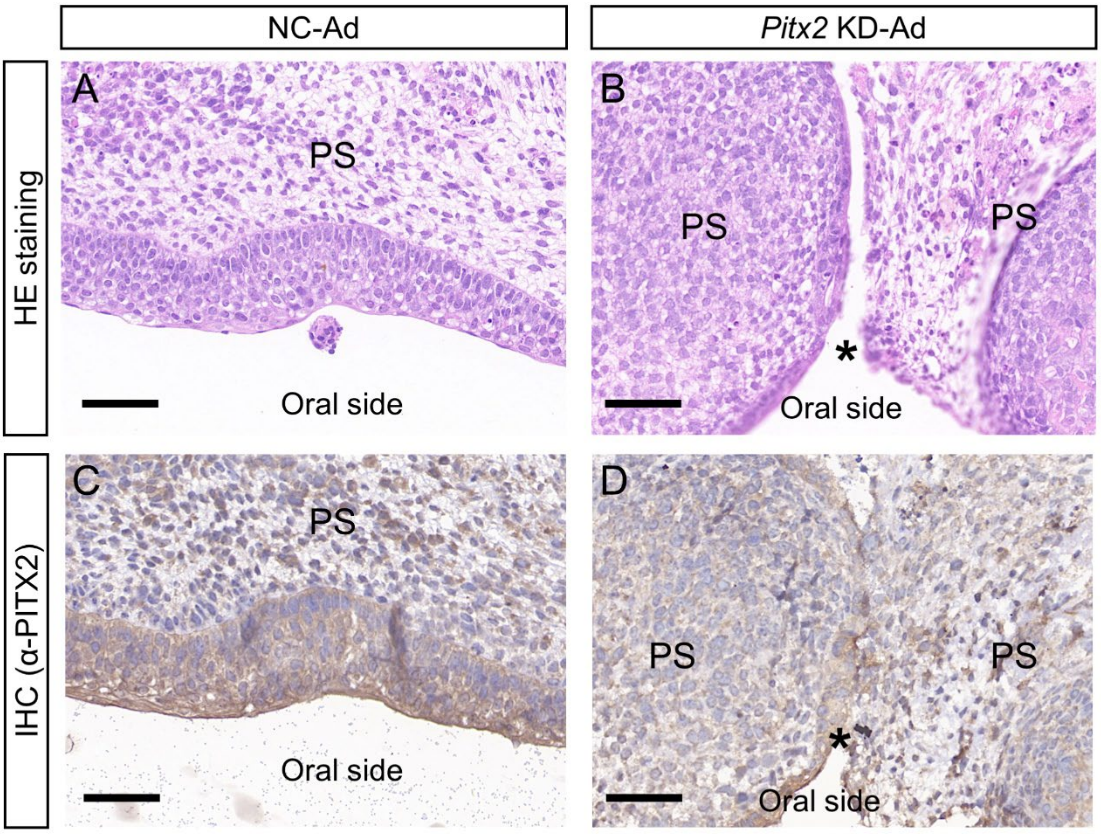


**Figure S16. PITX2 knockdown impairs adhesion of mouse palatal shelves *ex vivo*. (A and B)** The state of palatal adhesion was evaluated using HE staining after 72 hours of palatal organ culture. Samples were treated with **(A)** the negative control adenovirus (NC-Ad) or **(B)** *Pitx2* knock-down adenovirus (*Pitx2* KD-Ad). **(C and D)** IHC assays were conducted to detect PITX2 expression levels in palatal shelves after 72 hours of culture. Samples were treated with **(C)** NC-Ad or **(D)** *Pitx2* KD-Ad. In the *Pitx2* KD-Ad group, PITX2 expression in the periderm was reduced compared to that in the NC-Ad group. Asterisks indicate non-adhesion of palatal shelves. Scale bars, 50 μm. PS, palatal shelf.

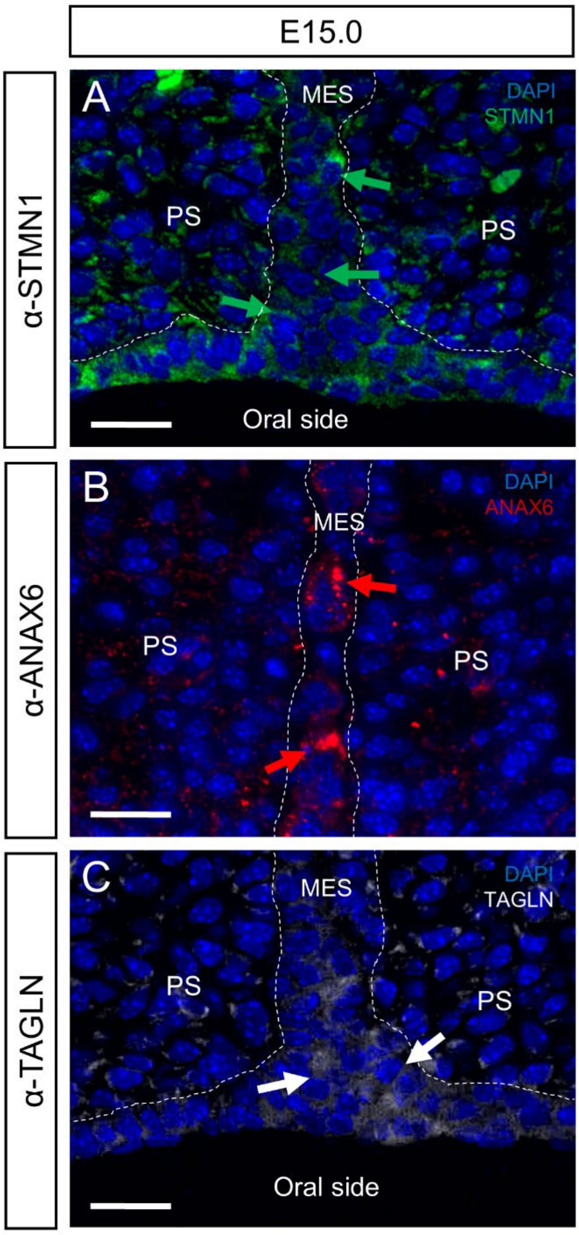


**Figure S17. Immunofluorescence validation of EMT, apoptosis, and migration contributing to the degeneration of periderm cells in the medial epithelial seam.** Coronal sections of E15.0 mouse embryos were analyzed using immunofluorescence assays for **(A)** STMN1, **(B)** ANXA6, and **(C)** TAGLN. Our analysis revealed that STMN1 (an EMT marker) and ANXA6 (an apoptosis marker) were expressed within the medial epithelial seam (MES) region. TAGLN, a marker for cell migration, was expressed in the region abutting the oral side. The expression patterns of these markers were consistent with the results obtained from the RNAscope ISH assay for *Vim*, *Igfbp3*, and *Csrp1* (refer to Figure 7), confirming that EMT, apoptosis, and migration collectively contribute to the degeneration of periderm cells in the MES. White dashed lines indicate the MES region and arrows point to the STMN1-, ANXA6-, and TAGLN-positive cells. Nuclei are stained with DAPI. Scale bars, 20 μm. PS, palatal shelf; MES, medial epithelial seam.

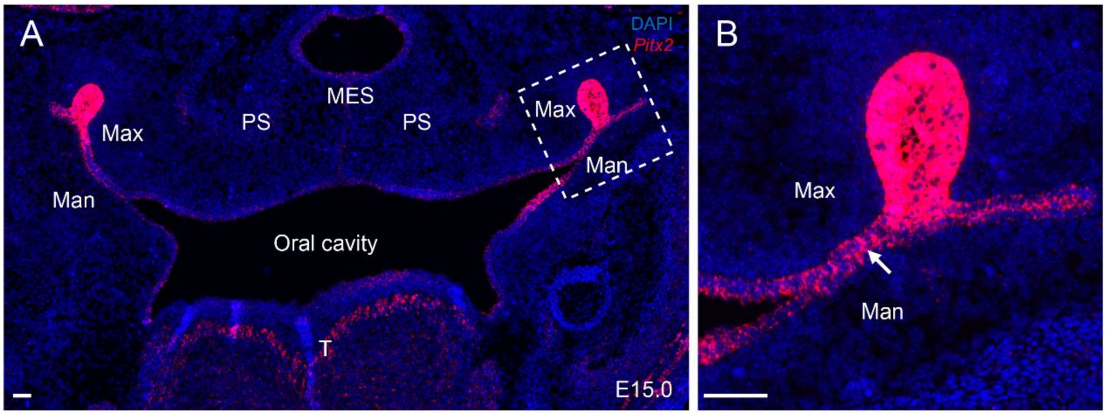


**Figure S18. RNAscope *in situ* hybridization (ISH) demonstrating the expression of *Pitx2* in the MES as well as widely in the oral epithelium.**

*Pitx2* was strongly expressed at the junction site between maxillary and mandibular tissue, where the two layers of epithelial cells are merging into a single layer (white arrow in B). Note that the expression of *Pitx2* does not lead to widespread pathological adhesion in the oral region. Dashed boxed areas are shown at higher magnification in B. Scale bars, 50 μm. T, tongue; PS, palatal shelf; MES, medial epithelial seam; Max, maxillary tissue; Man, mandibular tissue.

**Table S1 Summary of cell clusters during mouse palatogenesis**

| Cluster | Marker genes | Number of cells | | | | | Putative identity |
|---|---|---|---|---|---|---|---|
| | | E10.5 (N=12368) | E13.5 (N=8950) | E15 (N=9173) | E16.5 (N=10928) | Overall (N=41419) | |
| C0 | *Pax9, Barx1* | 33 (0.3%) | 1313 (14.7%) | 1145 (12.5%) | 1361 (12.5%) | 3852 (9.3%) | Developing palatal mesenchymal cells-posterior region |
| C1 | *Pax9, Barx1* | 25 (0.2%) | 872 (9.7%) | 889 (9.7%) | 1520 (13.9%) | 3306 (8.0%) | Developing palatal mesenchymal cells-posterior region |
| C2 | *Sp7, Alpl* | 43 (0.3%) | 1092 (12.2%) | 957 (10.4%) | 1172 (10.7%) | 3264 (7.9%) | Osteocyte lineage |
| C3 | *Bmp4, Msx1* | 57 (0.5%) | 588 (6.6%) | 1278 (13.9%) | 1082 (9.9%) | 3005 (7.3%) | Developing palatal mesenchymal cells-anterior region |
| C4 | *Cks2, Birc5* | 1017 (8.2%) | 704 (7.9%) | 527 (5.7%) | 562 (5.1%) | 2810 (6.8%) | Proliferating mesenchymal cells |
| C5 | *Sox2* | 1501 (12.1%) | 97 (1.1%) | 129 (1.4%) | 90 (0.8%) | 1817 (4.4%) | Epithelial progenitor cells |
| C6 | *Krt6a, Krt6b, Lypd3* | 12 (0.1%) | 628 (7.0%) | 600 (6.5%) | 285 (2.6%) | 1525 (3.7%) | *Krt6*+ cells |
| C7 | *Sox2* | 386 (3.1%) | 246 (2.7%) | 270 (2.9%) | 309 (2.8%) | 1211 (2.9%) | Epithelial progenitor cells |
| C8 | *Alas2, Hba-a1* | 25 (0.2%) | 131 (1.5%) | 209 (2.3%) | 379 (3.5%) | 744 (1.8%) | Red blood cells |
| C9 | *Crabp1* | 0 (0%) | 120 (1.3%) | 82 (0.9%) | 57 (0.5%) | 259 (0.6%) | Early-stage mesenchymal cells |
| C10 | *Col2a1, Col9a1* | 0 (0%) | 81 (0.9%) | 100 (1.1%) | 69 (0.6%) | 250 (0.6%) | Chondrocytes |
| C11 | *Elavl3* | 13 (0.1%) | 27 (0.3%) | 43 (0.5%) | 127 (1.2%) | 210 (0.5%) | Neuronal cell lineage |
| C12 | *Alas2, Hba-a1* | 2630 (21.3%) | 58 (0.6%) | 23 (0.3%) | 20 (0.2%) | 2731 (6.6%) | Red blood cells |
| C13 | *Trpm1, Six6, Aldh1a3* | 387 (3.1%) | 447 (5.0%) | 628 (6.8%) | 971 (8.9%) | 2433 (5.9%) | Eyes-related epithelial cells |
| C14 | - | 1834 (14.8%) | 72 (0.8%) | 29 (0.3%) | 34 (0.3%) | 1969 (4.8%) | Ambiguous cell type |
| C15 | *Egfl7, Cdh5* | 1031 (8.3%) | 56 (0.6%) | 41 (0.4%) | 91 (0.8%) | 1219 (2.9%) | Endothelial cells |
| C16 | *Alas2, Hba-a1* | 420 (3.4%) | 113 (1.3%) | 75 (0.8%) | 134 (1.2%) | 742 (1.8%) | Red blood cells |

| | | | | | | | |
|---|---|---|---|---|---|---|---|
| C17 | *Plp1, Mpz* | 56 (0.5%) | 91 (1.0%) | 132 (1.4%) | 202 (1.8%) | 481 (1.2%) | Schwann cells |
| C18 | *Myod1, Myog* | 11 (0.1%) | 120 (1.3%) | 37 (0.4%) | 20 (0.2%) | 188 (0.5%) | Myogenic precursor cells |
| C19 | *Trpm1, Six6, Aldh1a3* | 0 (0%) | 0 (0%) | 1 (0.0%) | 83 (0.8%) | 84 (0.2%) | Eyes-related epithelial cells |
| C20 | *Alas2, Hba-a1* | 56 (0.5%) | 470 (5.3%) | 661 (7.2%) | 688 (6.3%) | 1875 (4.5%) | Red blood cells |
| C21 | *Cd74, Rac2* | 36 (0.3%) | 495 (5.5%) | 392 (4.3%) | 350 (3.2%) | 1273 (3.1%) | Hematopoietic progenitor cells |
| C22 | *Lypd2, Cbr2* | 659 (5.3%) | 109 (1.2%) | 124 (1.4%) | 90 (0.8%) | 982 (2.4%) | Nasal epithelial cells |
| C23 | *Elavl3* | 639 (5.2%) | 19 (0.2%) | 27 (0.3%) | 23 (0.2%) | 708 (1.7%) | Neuronal cell lineage |
| C24 | *Sp7, Alpl* | 863 (7.0%) | 189 (2.1%) | 116 (1.3%) | 213 (1.9%) | 1381 (3.3%) | Osteocyte lineage |
| C25 | *Bmp4, Msx1* | 188 (1.5%) | 161 (1.8%) | 62 (0.7%) | 48 (0.4%) | 459 (1.1%) | Developing palatal mesenchymal cells-anterior region |
| C26 | *Myl9* | 304 (2.5%) | 282 (3.2%) | 181 (2.0%) | 300 (2.7%) | 1067 (2.6%) | Smooth muscle cells |
| C27 | *Krt14* | 26 (0.2%) | 281 (3.1%) | 209 (2.3%) | 433 (4.0%) | 949 (2.3%) | Epithelial basal cells |
| C28 | *Ambn, Amelx* | 116 (0.9%) | 88 (1.0%) | 206 (2.2%) | 215 (2.0%) | 625 (1.5%) | Dental cells |

*Note*: The percentages in parentheses indicate the proportion of each cluster at the corresponding time point.

**Table S4. Antibodies used in immunofluorescence assay for marker genes.**

| Marker | Cluster (s) | Cell type | Company | Cat.No. | Dilution | Reference |
|---|---|---|---|---|---|---|
| COL3A1 | C0-C4, C9, C10, C14, C18, C24, C25, C26 | Mesenchymal cells | Proteintech | 22734-1-AP | 1:100 | *Arthritis Res Ther* 2014[15] |
| TRP63 | C5-C7, C13, C19, C22, C27-28 | Epithelial cells | Proteintech | 12143-1-AP | 1:800 | *Sci Adv* 2019[16] |
| KRT6A | C6 | *Krt6*+ cells | Proteintech | 10590-1-AP | 1:200 | *Mech Dev* 2001[37] |
| KRT10 | C6.5.0 | Keratinized periderm cells I | Proteintech | 18343-1-AP | 1:100 | *J Allergy Clin Immunol* 2020 [46] |
| KLF4 | C6.5.1 | Keratinized periderm cells II | Proteintech | 11880-1-AP | 1:100 | *Sci Adv* 2019 [16] |
| IGFBP3 | C6.5.2 | Medial edge periderm cells | Proteintech | 10189-2-AP | 1:100 | *J Biol Chem* 1997 [55] |
| CLDN3 | C6.5.3 | Primitive periderm cells | Proteintech | 16456-1-AP | 1:100 | *Development* 2019[14] |
| ARHGAP29 | C6.5.3 | Primitive periderm cells | Proteintech | 12583-1-AP | 1:100 | *J Dent Res* 2017[44] |
| STMN1 | C6.5.2 | Medial edge periderm cells | CST | 3352 | 1:100 | *Cell Death Dis* 2022[58] |
| ANXA6 | C6.5.2 | Medial edge periderm cells | Proteintech | 12542-1-AP | 1:100 | *Clin Transl Med* 2020 [59] |
| TAGLN | C6.5.2 | Medial edge periderm cells | CST | 40471 | 1:100 | *Cell Death Dis* 2016 [60] |
